# Molecular mechanisms underlying single nucleotide polymorphism-induced reactivity decrease in CYP2D6

**DOI:** 10.1101/2024.01.04.574226

**Authors:** Daniel Becker, Prassad V. Bharatam, Holger Gohlke

## Abstract

Cytochrome P450 2D6 (CYP2D6) is one of the most important enzymes involved in drug metabolism. Genetic polymorphism can influence drug metabolism by CYP2D6 such that a therapy is seriously affected by under- or overdosing of drugs. However, a general explanation at the atomistic level for poor activity is missing so far. Here we show for the 20 most common single nucleotide polymorphisms (SNPs) of CYP2D6 that poor metabolism is driven by four mechanisms. We found in extensive all-atom molecular dynamics simulations that the rigidity of the I-helix (central helix), distance between central phenylalanines (stabilizing bound substrate), availability of basic residues on the surface of CYP2D6 (binding of Cytochrome P450 reductase), and position of arginine 132 (electron transfer to heme) are essential for an extensive function of the enzyme. These results were applied to SNPs with unknown effects and potential SNPs that may lead to poor drug metabolism were identified. The revealed molecular mechanisms might be important for other drug-metabolizing Cytochrome P450 enzymes.

**Table of Contents Graphic:** 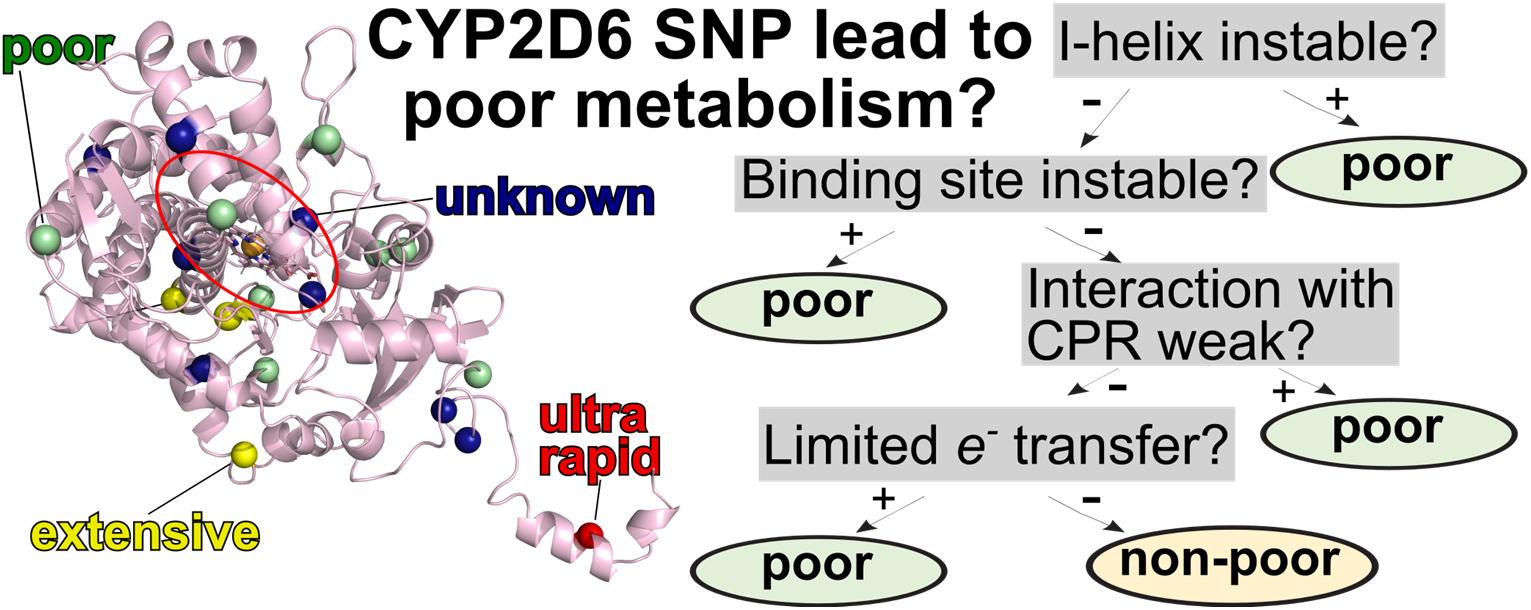

## Introduction

The isoenzyme cytochrome P450 (CYP) 2D6 plays a crucial role in human drug metabolism.^1^ It catalyzes the oxidation of especially lipophilic drugs that takes place as phase-1 biotransformation before excretion. Already in 1977, Mahgoub *et al.* described that differences in the hydroxylation rate of debrisoquine, an antihypertensive drug, depend on a single autosomal gene,^2^ called later CYP2D6 according to its position in the CYP gene superfamily.^3^ Since then, CYP2D6 polymorphism has raised increasing awareness during the development of small-molecule drugs because the polymorphism can lead to underdosing – leading to a failure of therapy or intoxication by drug metabolites – or overdosing – leading to intoxication by the drug itself.^4^

Differential mechanisms drive the differences in CYP2D6-related metabolism. The "very extensive" or "ultra-metabolizer" phenotype is mostly driven by gene duplication.^5^ This duplication results in the overexpression of the CYP2D6 enzyme. Around 7% of Caucasians show an "ultra-metabolizer" phenotype.^6^ Although an explanation at the genetic level for the "ultra-metabolizer" phenotype is often appropriate, mutations in the CYP2D6 gene can also lead to higher enzyme activity.^7^ The "poor-metabolizer" and "non-metabolizer" phenotypes are established either at the genetic level, probably related to regulatory factors^8^, or protein level, in which a combination of single nucleotide polymorphisms (SNPs) changes the enzymatic activity^9^. CYP2D6 is a highly polymorphic protein: So far, 165 genetic variants (alleles) have been discovered.^10^ Deciphering the molecular mechanisms underlying the SNP-induced reactivity decrease of CYP2D6 is of utmost interest from a fundamental and drug development point of view.

CYP2D6, like all human CYP enzymes, is a membrane-associated and heme-containing protein, where the heme is buried and only accessible via channels.^11, 12^ The oxidation reaction catalyzed by CYP enzymes occurs in multiple steps, starting from a heme with Fe(III) configuration (Figure 1A):^13^ First, the ligand usually binds to the heme iron by displacing a distal water molecule. Afterward, an electron is transferred to the heme from Cytochrome P450 Reductase (CPR), a membrane-bound enzyme required for electron transfer from NADPH to CYP.^14, 15^ In mammalian CYPs, the electron is transferred with arginine (in CYP2D6: Arg132), acting as a bridge between the flavine mononucleotide (FMN) of CPR and the heme iron.^16^ Fe(II) of heme now binds an oxygen molecule, and a second electron is transferred, yielding - after protonation - a ferric hydroperoxyl complex that rapidly separates one water molecule and results in a ferryl-coupled porphyrin radical cation. Finally, the highly reactive radical cation oxidizes the substrate, and after the product egress, the cycle is completed.^17^

**Figure 1:**
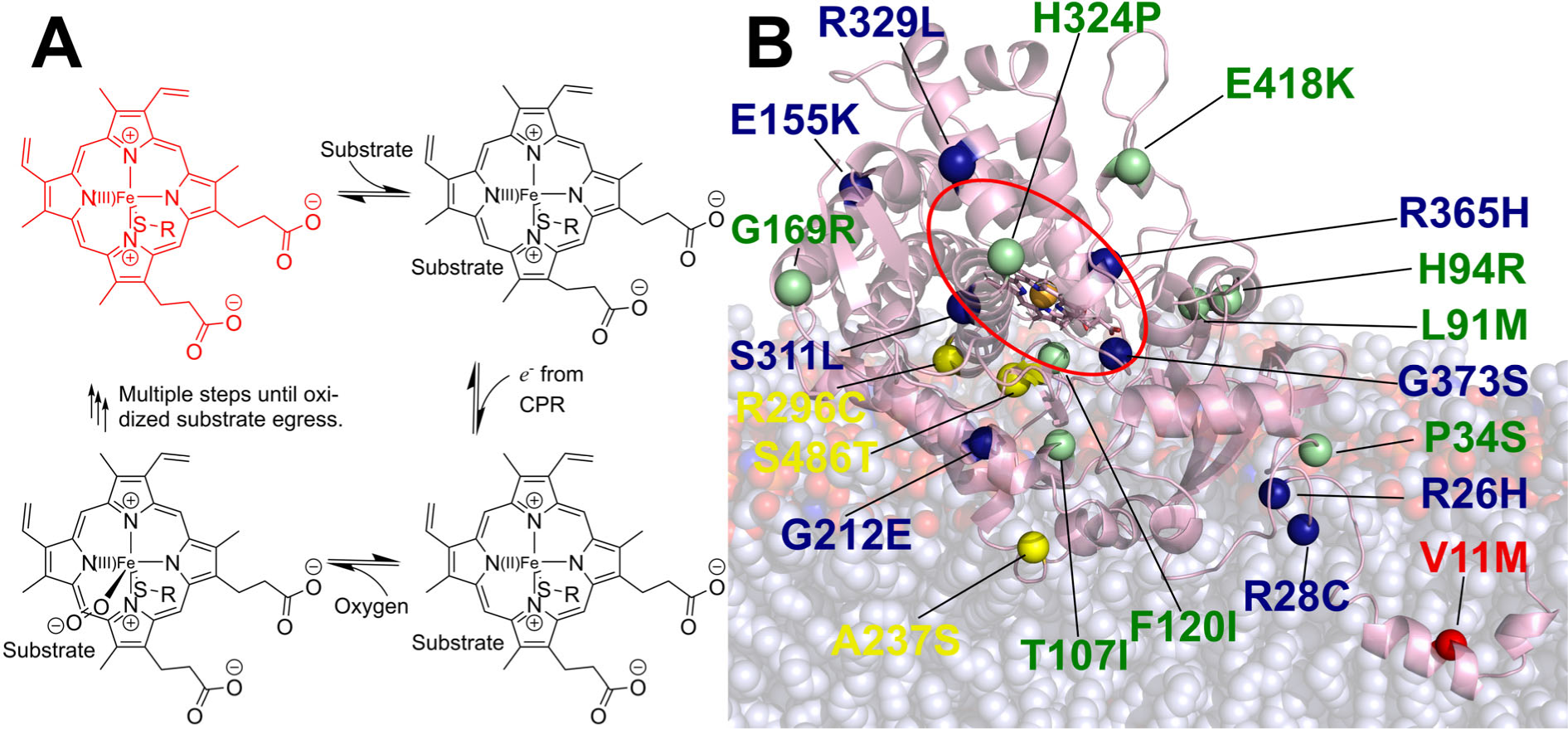
CYP2D6 is a heme-containing, membrane-associated enzyme. **A:** Parts of the reaction cycle of CYP enzymes underline the importance of substrate binding as well as electron transfer from CPR. The scheme was adapted from ref.^13^. **B:** Overview of the 20 most frequent SNPs mapped onto CYP2D6 (pink). Mutations that lead to poor metabolism are colored in green, variants with no effect are colored in yellow, V11M leading to an ultra-rapid metabolization is colored in red, and variants with unknown effects are colored in blue. Cα atoms of the positions are shown as spheres. The orange sphere in the center indicates the heme iron in the center of the porphyrin ring, which is shown as sticks.

CPR undergoes a complex mechanism consisting of open and closed states to provide electrons to CYP. First, the closing of the FMN domain establishes an interaction between this domain and the flavine dinucleotide (FAD) domain.^18^ Due to this interaction, electrons of FAD, originating from NADPH, are transferred to FMN. Second, CPR reopens and forms a complex with CYP, mainly driven by polar interactions. The surface of the CYP enzyme is positively charged in the interaction region by several arginine and lysine residues (in the case of CYP2D6: Arg129, Arg133, Arg140, Lys146, Lys429, Arg440, and Arg450), while the surface of the FMN domain is negatively charged due to aspartic acid and glutamic acid residues.^19^

Effects on CYP2D6 reactivity related to different alleles, including combinations of mutations, were investigated previously in wet lab experiments and computational studies.^9, 20–22^ Due to the many naturally occurring alleles, only investigations of a subgroup of alleles have been possible. In these studies, different aspects of how mutations can affect the catalysis steps have been pointed out. I) Decreased activity can be correlated with an instability of the expressed enzyme variant.^23^ II) Mutations of Phe120 and Phe483 affect the substrate binding.^24–26^ III) As to the arginines and lysines playing a crucial role in the interaction between CYP and CPR, the allele CYP2D6*31 (combining SNPs R296C, R440H, and S486T; listed in the Pharmacogene Variation Consortium database) has a mutation in one of these positions, likely underlying that this CYP2D6 variant has no enzymatic activity *in vivo* and *in vitro*.^27^ IV) The bridging arginine between FMN of CPR that supports the electron transfer ^16^ is another crucial residue; as a substitution probably prevents electron transfer to the heme, i.e., leads to inactivity, no substitution has been documented.

In this work, we aim to elucidate the effect of mutations found in naturally occurring alleles on CYP2D6 activity on an atomistic level. Due to the large number of natural mutants, our study focuses on the twenty most common SNPs in the global population (Figure 1B). We performed extensive all-atom MD simulations with a cumulated simulation time of over 200 µs. On this basis, we pursue a mechanism- and hypothesis-driven evaluation to detect if and how an SNP probably leads to a decreased reactivity of the mutant.

## Materials and methods

### Structural models of CYP2D6 mutants

We used the equilibrated structural model of CYP2D6 wildtype generated in our previous work by modeling the globular part and transmembrane helix separately with TopModel^28^ and docking both structures with spatial restraints as a basis^29^. Repeating the modeling of CYP2D6 wildtype by AlphaFold v2^30^ yields a root-mean-square deviation (RMSD) < 2 Å, confirming the accuracy of our previous modeling. The selected mutations were introduced into the structural model using PyMOL, thereby paying attention that the rotamer with the least clashes was chosen.^31^ The protonation state of the protein residues was estimated for pH 7.4 with PROPKA3^32^ using HTMD.^33^ The generated models were embedded by PACKMOL-Memgen^34^ into a bilayer membrane with a composition reflecting the main lipid components of the human endoplasmic reticulum^35^ (CHL:DOPC:DSPC:DAPC:DOPE 10:22:13:19:21). Potassium chloride was added with 0.15 M concentration. The orientation of the structural model in the membrane was determined with MEMEMBED.^36^

### Molecular dynamics simulations

The GPU particle mesh Ewald implementation^37^ of the AMBER18^38^ molecular simulations suite was used with ff14SB parameters^39^ for the protein, Lipid17 parameters^40^ for the membrane, and OPC as the water model^41^ and Li/Merz parameters for the ions^42^. Parameters for the heme and cystine residue forming the S-Fe bridge between heme and the protein were taken from Shahrokh *et al.*^43^. Because covalent bonds to hydrogens were constrained with the SHAKE algorithm^44^, a time step of 2 fs was used. The cutoff for nonbonded interactions was set to 10 Å.

Production runs were prepared as described by Schott-Verdugo et al.^45^, but the minimization without restraints was performed for 40,000 steps. Langevin dynamics^46^ with a friction coefficient of 1 ps^-1^ was used to keep the temperature at 310 K. Initially, the minimized systems were heated by gradually increasing the temperature from 0 K to 310 K for 280 ps under NVT conditions. The system density was adjusted using NPT conditions at 1 bar for 2 ns. Production runs were then performed using the same NPT conditions. The pressure was controlled with the Berendsen barostat^47^ with semi-isotropic pressure scaling, coupling the membrane (x,y) plane.

Geometric analyses were performed with cpptraj^48^ from the AmberTools suite^49^.

### Conformational sampling

To improve the robustness of the analyses and quantify the statistical uncertainty of the results, we carried out all analyses on ensembles of conformations generated from ten MD trajectories of 1 µs length for each of the enzymes. The independence of the replicas was ensured by using heating processes with a different random seed for each replica. The structures remain structurally stable with a backbone RMSD < 3.5 Å compared to the starting structure, considering the protein part without the transmembrane helix, which moves independently from the globular part.

### Constraint network analysis

We analyzed static properties, i.e., structural rigidity and its opposite flexibility,^50^ of the 20 CYP2D6 mutants and the wildtype. Therefore, the enzymes were represented as constraint networks, where atoms are the nodes, and covalent and noncovalent bonds constitute constraints in between.^51^ Noncovalent interactions such as hydrogen bonds, salt bridges, hydrophobic tethers, and stacking interactions contribute most to biomolecular stability. Using an empirical energy function allows for quantifying the strength of hydrogen bonds and salt bridges.^52^ By gradually removing these polar noncovalent constraints from an initial network representation of a biomolecule according to a cutoff energy *E*_cut_, a succession of network states *σ* is generated that forms a ’constraint dilution trajectory’.^53, 54^ To this end, hydrogen bonds and salt bridges are removed in the order of increasing strength such that for network state *σ*, only those hydrogen bonds are kept that have an energy *E*_HB_ ≤ *E*_cut_(*σ*). Performing rigidity analysis^55^ on such a trajectory reveals a hierarchy of structural stability that reflects the modular structure of biomolecules in terms of secondary, tertiary, and super-tertiary structure.

By this, we obtained for each system a neighbor stability map (*rc_ij_*_,neighbor_(*E*_cut_(*σ*))) that contains information accumulated over all network states *σ* along the trajectory ^56, 57^ in that it monitors the persistence of rigid contacts for pairs of residues during a constraint dilution process. From the stability maps, we calculated *E_i,_*_CNA_ (eq. 1), the sum of energies associated with rigid contacts between residue *i* to all other residues.

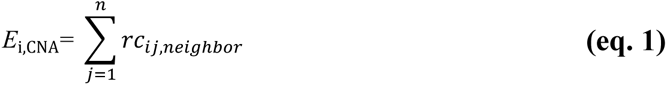

We also calculated *Ē*_region,CNA_ (eq. 2), where "region" is either the entire protein, the F/G region, or the I-helix, as the average of the chemical potential of rigid contacts between the *n* residues within a region.

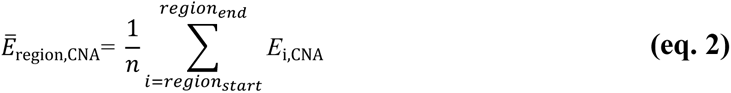

The computations were done with the Constraint Network Analysis (CNA) program (version 4.0) developed by us.^54^ For further details, see our previous work^29^.

### Generation of a structural model of the CYP2D6-CPR complex

The structural model of the complex of CYP2D6 and the FMN domain of CPR (Uniprot ID: P16435) was obtained through protein-protein docking using the HADDOCK webserver with default settings^58, 59^ and is used for visualization purposes. For the FMN domain, residues 142, 144, 147, 179, and 209 were considered involved in the interaction. For CYP2D6, residues 129, 133, 140, 146, 429, and 440 were considered involved. The involved residues were chosen based on literature^15^ and serve as putative interacting residues during the docking.

## Results

### Identification of relevant SNPs

The most common SNPs were identified based on the Uniprot entry (Uniprot-ID: P10635)^60^ that reports on 50 SNPs. We restricted ourselves to the 20 most frequent mutations with a relative frequency of at least 10^-4^ in the global population. This frequency was taken from the dbSNP database.^61^ Data from ClinVar,^62^ PharmVar,^27^ and the Human Gene Mutation Databank (HGMD)^63^ were used to assess the clinical relevance of the mutations (see Table 1). For variants leading to decreased reactivity, we used the term "poor", for the wildtype and variants with no change in reactivity, the term "extensive" was used, the variant associated with ultrarapid metabolism was labeled as "ultrarapid", and the variants with unknown effects on reactivity were labeled as "unknown".

**Table 1:** Classification of clinical effects of CYP2D6 SNPs investigated here and source of the classification.

| SNP | ClinVar<sup>62</sup> | HGMD<sup>63</sup> | PharmVar<sup>27</sup> | Class |
| --- | --- | --- | --- | --- |
| WT |  |  |  | extensive |
| V11M | likely benign | ultrarapid metabolizer |  | ultrarapid |
| A237S | likely benign |  | normal | extensive |
| R296C | likely benign |  |  | extensive |
| S486T | likely benign |  | normal | extensive |
| P34S | poor metabolism | poor metabolizer |  | poor |
| L91M |  | poor metabolizer |  | poor |
| H94R |  | poor metabolizer |  | poor |
| T107I | likely benign | poor metabolizer |  | poor |
| F120I |  | poor metabolizer |  | poor |
| G169R | poor metabolism | poor metabolizer |  | poor |
| H324P | likely benign | poor metabolizer | no function | poor |
| E418K |  | reduced enzyme activity |  | poor |
| R26H |  |  |  | unknown |
| R28C |  |  |  | unknown |
| E155K |  |  |  | unknown |
| G212E |  |  |  | unknown |
| S311L |  |  |  | unknown |
| R329L |  |  |  | unknown |
| G373S |  |  |  | unknown |
| R365H |  |  |  | unknown |

### General strategy to identify atomistic mechanisms

For this study, we assumed that mutations documented as extensive and ultrarapid can be used as a comparison group (CG) to identify the atomistic mechanisms leading to decreased activity in the other variants. The variants were investigated as to different aspects relevant to the catalyzed reaction, i.e., enzyme stability, substrate binding, binding of CPR, electron transfer, and substrate access as detailed below. We then defined criteria for determining whether an aspect is significantly different from the CG, which could also be applied to characterize other variants not included in this dataset.

To do this, for a mechanism *A*, we identified the upper or lower limit (depending on the mechanistic relation) of a criterion *u*_A,limit_ (eq. 3) that shows a significant difference for a given significance level *p* with the average criterion of the control group *ū_A_*_,CG_ with standard deviation *σ_A_*_,*CG*_ (eq. 4). To obtain the most robust *u*_A,limit_ estimate according to our dataset, we respectively considered the largest standard deviation *σ_A_*_,*nonCG*,*max*_ incurred for any of the variants that are not in the control group (nonCG).

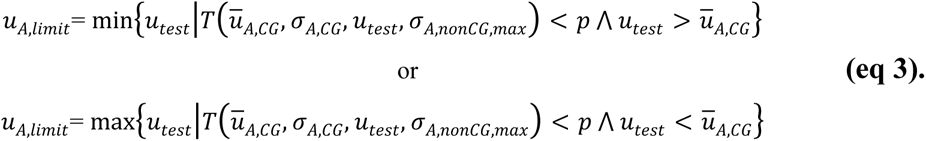

where *p* was set to 0.05 and *T* is the *t*-test function implemented in SciPy v1.7.3.^64^

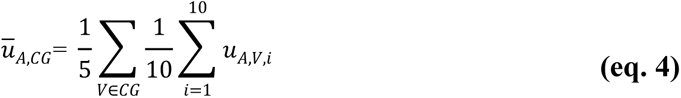

### Structural instability of the I-helix

The first molecular mechanism relates to a loss of rigid contacts from the I-helix (residues L293-L323) to other parts of CYP2D6 as deduced from CNA. The I-helix is the central and one of the least mobile helices within the enzyme (Figure 2A, S1A,B). The loss of rigid contacts is indicated by a decreased average chemical potential (eq. 2) in this region, which is significant only for the substitution H324P with respect to CG and all other nonCG (*p* < 0.01, two-sided *t*-test) (Figure 2B).

**Figure 2:**
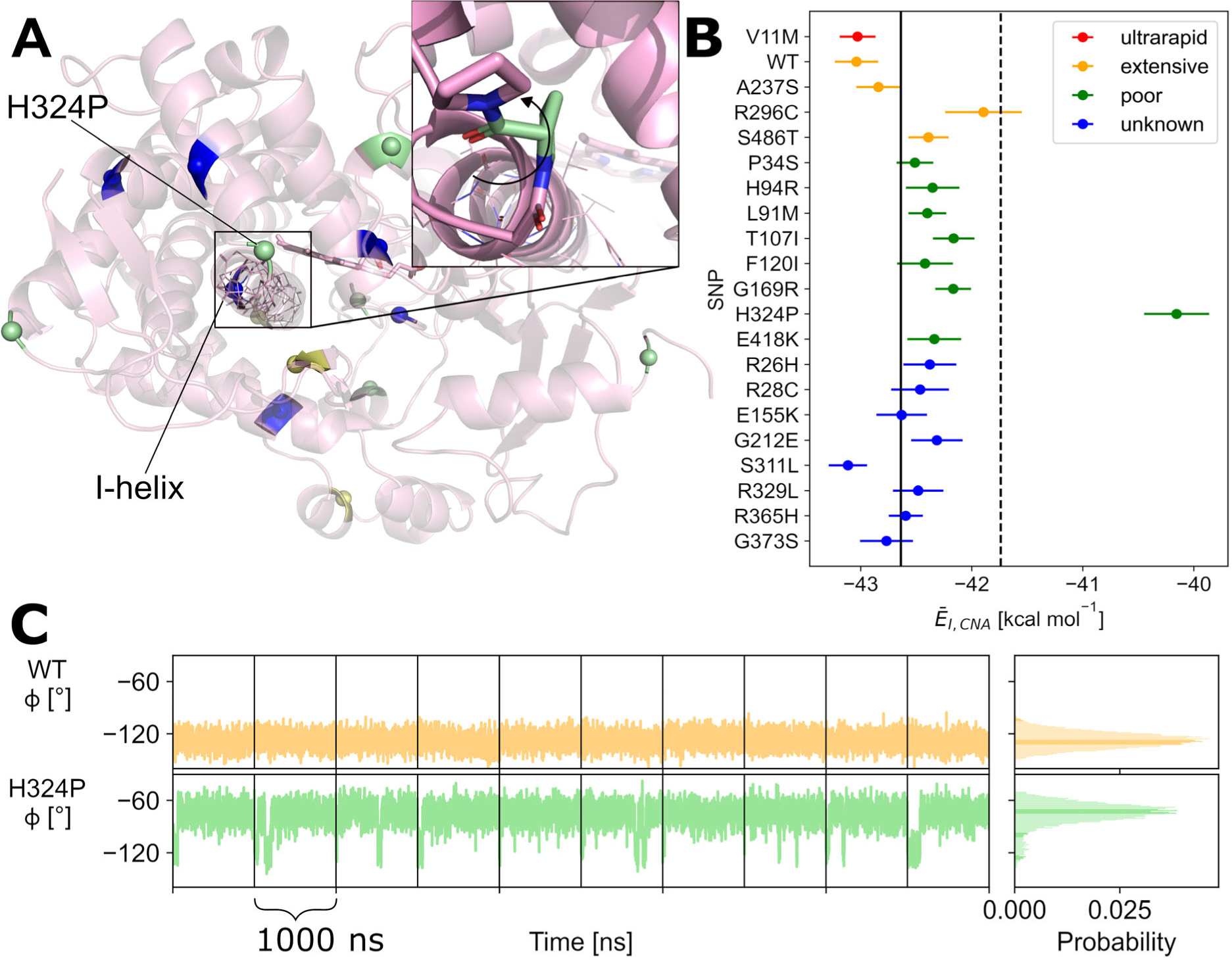
Structural instability of the I-helix leads to poor metabolism of CYP2D6 H324P. **A:** Structure of the globular part of CYP2D6 (pink) shown with the I-helix marked with backbone lines and positions of SNPs as shown in Figure 1. The heme group is shown as sticks. In the blowup, the H324P variant is shown with the measured ϕ angle indicated. **B:** Average chemical potential of rigid contacts of I-helix residues (*Ē*_I,CNA_) shown for each investigated SNP. For all variants except H324P, the value is approximately between -43 and -42 kcal mol^-1^. Only H324P differs significantly (*p* < 0.01, two-sided t-test) from the comparison group, leading to a decreased structural stability of the I-helix. The continuous vertical line denotes the mean of *Ē_I,_*_CNA_ of the comparison group (eq. 2); the error bars denote the SEM determined by error propagation along 10 independent simulations. The dashed vertical line denotes the limit for a significantly different value (*p* < 0.01, two-sided *t*-test, eq. 3). **C:** The ϕ angle, i.e., dihedral angle C_i-1_-N_i_-Cα_i_ -C_i_, of residue 324 in the wildtype and the H324P variant is shown over the simulation time (left) and as frequency distribution (right). Vertical lines separate the 10 replicas. In each replica of H324P, a shift to ∼-75° is visible, while the dihedral remains at ∼ - 120° in all replicas of the wild type.

From a structural viewpoint, proline is known for disrupting the secondary structure,^65^ which may be intensified by the proline at position 325 already being present in the wild type. In the H324P variant, the ϕ dihedral angle (C_i-1_-N_i_-C*α*_i_-C_i_) of this residue changes to ∼ -75° (loop conformation) from ∼ -120° (β-sheet-like conformation) in the wildtype (Figure 2C). Due to this change, the loop between I-helix and J-helix is elongated by one residue (WT: L323-P325; H324P: L323-D326), which may contribute to destabilizing the I-helix. Furthermore, the replacement of the histidine sidechain leads to more space for the C-terminal loop of CYP2D6, which also contains C443 interacting with the heme group (Figure S2A,B,C). With H324, the C-terminal loop formed more rigid contacts to the I-helix than with a proline at this position, which led to the decrease in structural stability of the I-helix in case of the substitution (Figure 2B).

### Distortion of the binding site integrity

F120 and F483 are important in substrate binding, especially for aromatic substrates (Figure 3A).^24–26^ The two phenylalanines build a hydrophobic dome that narrows down the binding pocket and builds a hydrophobic environment close to the heme necessary for the correct binding of the substrate.^66^ In our MD simulations, we saw large and significant differences (*p* < 0.01, two-sided *t*-test) in the distance between both phenylalanines in the variants T107I and G169R compared to the CG (Figure 3B). This suggests that substrate binding could be disfavorably impacted.

**Figure 3:**
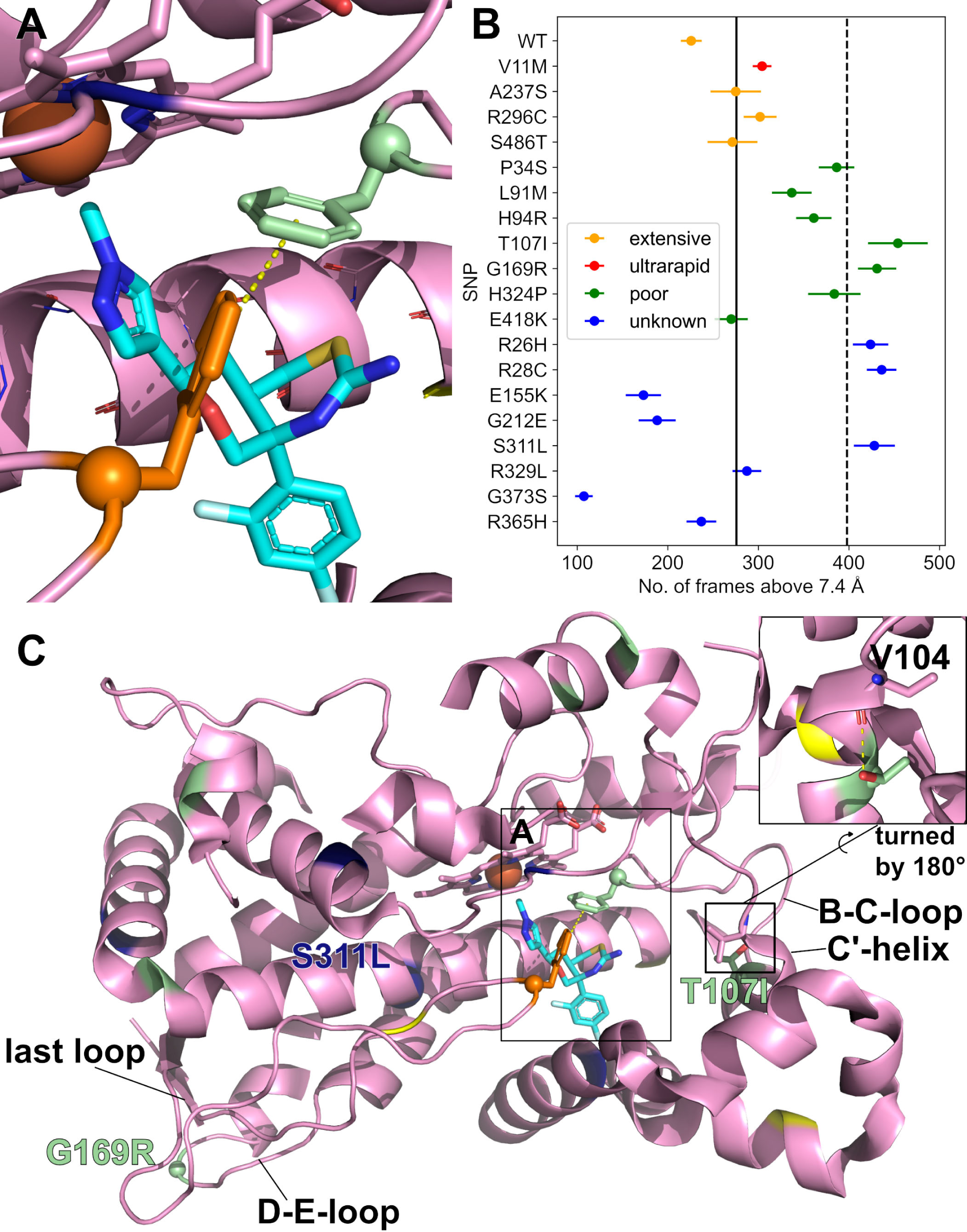
Importance of F120 and F483 in substrate binding. **A:** Zoom into the crystal structure of the active site of CYP2D6 (pink) binding inhibitor BACE-1 (cyan) (PDB-ID: 4XRY). The heme and ligand are shown as sticks; F120 is marked in green and F483 in orange. The iron within the heme is shown as a sphere. **B:** The number of frames with a distance between F120 and F483 above 7.4 Å is depicted for all investigated variants, except F120I. The distance was measured between the phenyl rings. The continuous vertical line denotes the mean distance between F120 and F483 of the comparison group (eq. 3); the error bars denote the SEM determined by error propagation along 10 independent simulations. The dashed vertical line denotes the limit for a significantly different value (*p* < 0.01, two-sided t-test, eq. 3). **C:** Crystal structure of CYP2D6 bound to BACE-1. Locations of mutations T107I and G169R are labeled. The influence on the distance between F120 and F483 could be explained by the vicinity of both mutations to the loops F120 (green) and F483 (orange) are part of. T107 is stabilizing the C’ helix via hydrogen bonding with the backbone oxygen of V104 shown in the blowup. Breaking this by an exchange of threonine by isoleucine leads to conformational changes of the B-C-loop, of which F120 is part. In the G169R variant, a small glycine is mutated to a voluminous arginine in the D-E-loop, which is close to the last loop in the sequence behind the L-helix, of which F483 is part.

For T107I, the change in the distance is probably caused by a conformational change in the C’-helix where a hydrogen bond from the threonine hydroxyl group to the V104 backbone oxygen cannot be established anymore. The missing interaction leads to a higher variance of the ψ dihedral angle (N_i_-C*α*_i_-C_i_-N_i+1_) at P105, with significantly more positive values in T107I than in the wildtype (Figure S3) (*p* < 0.01, two-sided *t*-test). The C’-helix is linked to the B-C-loop, which comprises F120 (Figure 3C).

G169R is part of the D-E-loop that interacts with the C-terminal loop by a hydrogen bond between F172 in the D-E-loop and L492 one residue before the β3-2-strand (Figure S4A,B,C). In the G169R variant, we observed a decreased distance between the C_α_ of R169 and the C_α_ of P496, which is the last residue of the β3-2-strand caused by electrostatic interactions of the guanidino group of R169 with the S168 hydroxyl group and the backbone nitrogen of P496 (Figure S4D-H). Due to the interaction between both loops, the shift in the D-E-loop is transferred to the C-terminal loop, which comprises F483.

In the F120I variant, the binding of aromatic substrates is likely hindered due to the missing π-interactions; at least one aromatic ring occurs in 99% of a database containing more than 3,500 compounds published by the medicinal chemistry departments of AstraZeneca, Pfizer, and GlaxoSmithKline.^67^ Thus, the variants T107I, G169R, and F120I are probably poor metabolizing enzymes due to changes in the substrate binding strength. If we apply the criterion to the group of unknown SNPs, R26H, R28C, and S311L lead to significantly higher distances between F120 and F483, which suggests that substrate binding is also decreased for these variants.

### Reduced interaction with Cytochrome P450 reductase

For the reaction cycle of CYP enzymes, the transfer of an electron from Cytochrome P450 reductase (CPR) is mandatory. CPR itself undergoes a complex cycle that requires binding and unbinding from the CYP enzyme.^68^ The binding between CYP and CPR is mediated by basic residues (positive charges) on the CYP side and acidic residues (negative charges) on the CPR side.^15, 69^ For CYP2D6, the basic residues are: R129, R133, R140, K146, K429, R440, and R450 (Figure 4A,B). For successful binding, the availability of these residues plays a crucial role. All of them do not form intramolecular interactions during the MD simulations except R450, which forms a salt bridge with E150. This salt bridge occurs in all investigated variants, but more often in L91M (significant: *p* < 0.01, t-test) and G373S (Figure 4C). This is also reflected by the average chemical potential of the rigid contact between R450 and E150 (*Ē*_R450-E150,CNA_; eq. 2) (Figure 4D). Hence, the formation of this salt bridge may lead to reduced binding of CYP2D6 L91M and CYP2D6 G373S to CPR and a lack of electrons for the metabolizing reaction.

**Figure 4:**
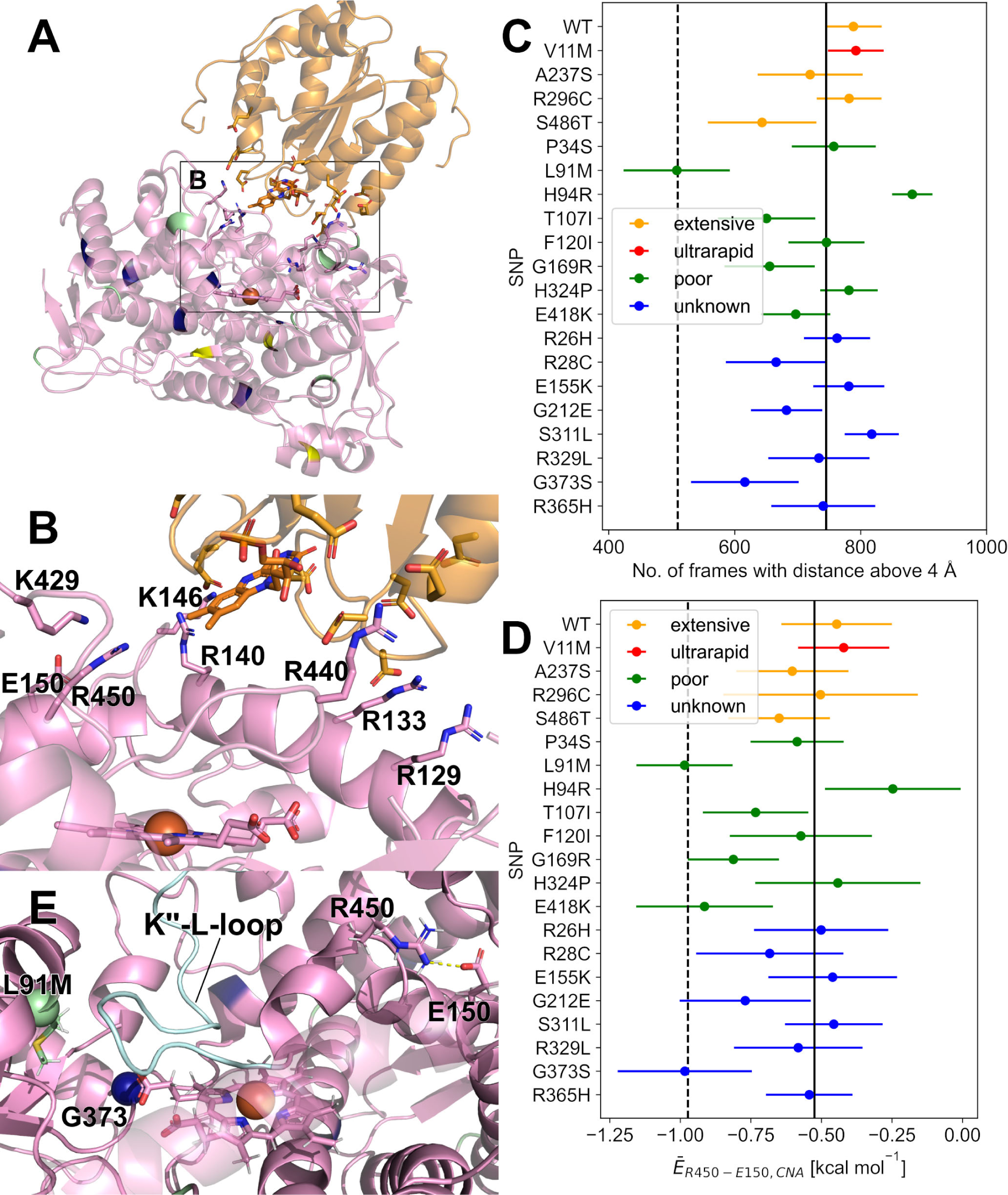
Arginine 450 plays an important role in CPR binding. **A:** Interface of CPR (yellow) and CYP2D6 (pink) derived by docking using the HADDOCK web server ^58^. E150 does not take part in the interaction with CPR but can form a salt bridge with R450. Heme and charged residues within the interface are shown as sticks. The iron within the heme is shown as a sphere. The residues are colored according to their effect on metabolism. **B:** The close-up view reveals the importance of charged residues for the interaction between CPR and CYP2D6. While the interface of CYP2D6 is mainly positively charged, the interface of CPR is negatively charged due to many glutamic and aspartic acids. **C:** The number of frames with a distance between R450 and E150 above 4 Å is depicted for all investigated variants. The distance was measured as the closest distance between all combinations of sidechain N’s of arginine and sidechain O’s of glutamic acid. Above 4 Å, the salt bridge is not considered formed^70^. Then, R450 is free to interact with CPR. The continuous vertical line denotes the mean distance between R450 and E150 of the CG; the error bars denote the SEM determined by error propagation along 10 independent simulations. The dashed vertical line denotes the limit for a significantly different value (*p* < 0.01, two-sided t-test). **D:** Average chemical potential of rigid contacts (*Ē*_R450-E150,CNA_) between R450 and E150. The salt bridge in the L91M and the G373S variants is significantly more stable than in all other variants. The continuous vertical line denotes the mean of *Ē*_R450-E150,CNA_ of the CG; the error bars denote the SEM determined by error propagation along 10 independent simulations. The dashed vertical line denotes the limit for a significantly different value (*p* < 0.01, two-sided t-test). **E:** The close-up view on CYP2D6 L91M shows that L91M but also G373S may influence the salt bridge stability between R450 and E150 by a shift of the K’’-L-loop (cyan) due to larger space requirements of the substitutions compared to the wildtype residues. The locations of L91M and G373 are shown as spheres. The M91 side chain is shown as sticks, as are R450 and E150.

L91M influences the salt bridge prevalence between R450 and E150 via the K’’-L-loop of CYP2D6 (Figure 4E). L91M is part of the B-helix, which is close to the K’’-L-loop. In MD simulations of the L91M variant, we observed a significantly decreased distance between the K’-L-loop (C_α_ of S437, G439, and R441) and the β1-3-strand (C_α_ of L395) compared to the CG, which is located below the B-helix (Figure S5A-E) (*p* < 0.01, two-sided *t*-test). The conformation of the K’’-L-loop itself does not change (Figure S5F). The movement of the K-L-loop leads to a change in the relative position of R450, which is close to the L-helix, which increases the prevalence of salt bridge formation with E150.

### Reduced electron transfer to heme mediated by R132

For the electron transfer from the flavin mononucleotide of CPR to the heme group of CYP, R132 of CYP, located between both cofactors, is needed (Figure 5A,B).^16^ R132 is part of the C-helix and is on average 7.8 Å away from the heme iron. For the three variants E418K, H94R, and P34S with decreased reactivity, we found that the distance between R132 and the iron of the heme group is on average significantly increased compared to the CG (Figure 5C). Thus, these variants are probably poor metabolizing enzymes due to the lower probability of an electron transfer via R132. The same criterion is fulfilled by R26H, R329L, and G373S of the unknown group.

**Figure 5:**
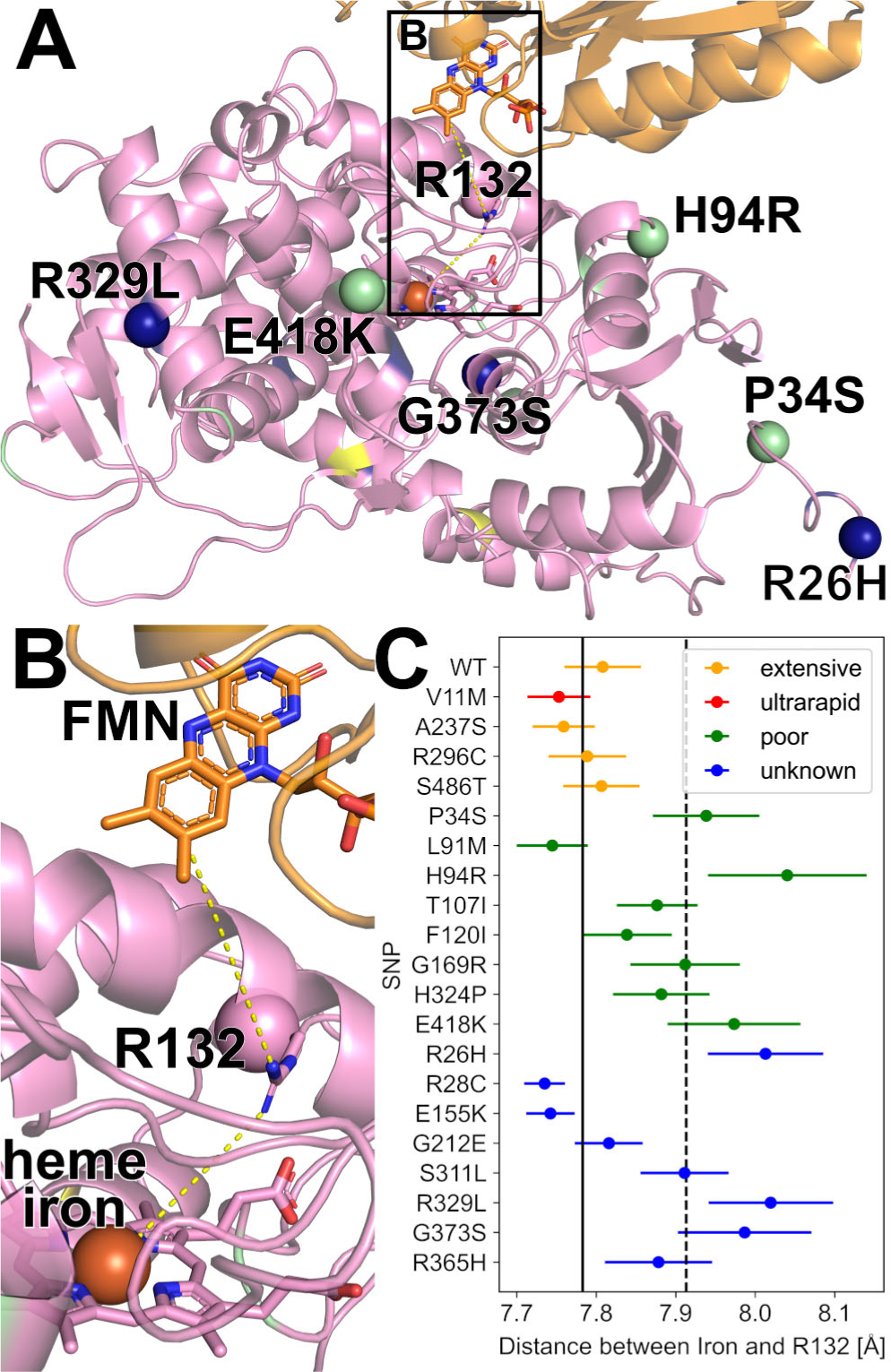
R132 plays a crucial role in electron transfer from CPR to CYP2D6. **A:** Interface of CPR (yellow) and CYP2D6 (pink) derived by docking using the HADDOCK web server ^58^. Heme and flavin mononucleotide are shown as sticks as is R132. The iron within the heme is shown as a sphere. All Cα-atoms of substitutions in variants that show a significantly higher distance between iron and R132 are shown as a sphere. **B:** Close-up view of the interface between CYP2D6 and CPR. The route of the electron transfer according to ref.^16^ is indicated with dashed yellow lines. Thus, the distance between R132 and the heme iron is an important factor for the likelihood of an electron transfer. **C:** Average distance between R132 and the heme iron. The continuous vertical line denotes the mean of the distance of the CG; the error bars denote the SEM determined by error propagation along 10 independent simulations. The dashed vertical line denotes the limit for a significantly different value (*p* < 0.01, two-sided *t*-test). The distance is measured as the closest distance between the terminal nitrogen atoms of arginine and iron.

E418K is part of the K’’-helix and at least 31.9 Å away from R132. Its sidechain points to the protein’s surface (Figure S6A). Our MD simulations revealed a minor conformational change at the K’’-helix, due to which the neighboring K429 is shifted towards the K’’-L-loop (Figure S6B). Consequently, the distance to A449 is significantly decreased by 0.2 Å (*p* < 0.01, two-sided *t*-test) (Figure S6C). This shift is then transmitted from A449 to E446 (Figure S6D) and from E446 to K140 (Figure S6E), which is part of the C-helix and has a constant distance to R132 (Figure S6F). These changes lead to a significantly higher distance between R132 and heme iron (Figure S6G).

H94R is part of the B-helix, which is spatially located between the β2-1-strand and the K’’-L-loop (Figure S7A-C). Our MD simulations revealed that the distance between the B-helix and the β2-1-strand (measured as the distance between Cα 94 and Cα 383 as the center of the β2-1-strand) is significantly increased by 0.7 Å (*p* < 0.01, two-sided *t*-test) (Figure S7D). Due to this shift, the interaction between the C-terminus of the B-helix and the C’-C-loop is affected, i.e., the hydrogen bond between E96 and Y124 is less often formed in H94R than the wildtype (*p* < 0.01, two-sided *t*-test) (Figure S7E). This leads to an overall higher displacement of the R132-containing C-helix in relation to the heme (Figure S7F).

P34S is part of the loop between the TM-helix and A-helix. Together with Y33, it stabilizes the β2-sheet by holding Y33 in between V68 (β1-1-strand) and F387 (β2-2-strand) (Figure S8A-C). In our MD simulations of P34S, we observed that Y33 moves away from this position (*p* < 0.01, two-sided *t*-test), which leads to a reorientation in the β2-2-strand, and the aromatic ring of F387 adopts the position of Y33 (Figure S8D). The consequence of the different conformation of the β2-2-strand is a shift of the B-helix, which interacts with the β2-2-strand as described above. The shift of the B-helix is transmitted via the interaction of E96 and Y124 to the C-helix, which contains R132.

### Unchanged stability of the F/G-Region

In our previous work, we showed that the structural stability of the F/G-region (residues P200 – M260), containing the F- and G-helices next to the substrate entrance channel (Figure 6A), determines the substrate promiscuity of drug-metabolizing human CYP enzymes.^29^ This is probably caused by the lid effect of this region. The stability of the F/G-region was determined by the average chemical potential due to rigid contacts in this region (*Ē*_FG,CNA_, eq. 2).

**Figure 6:**
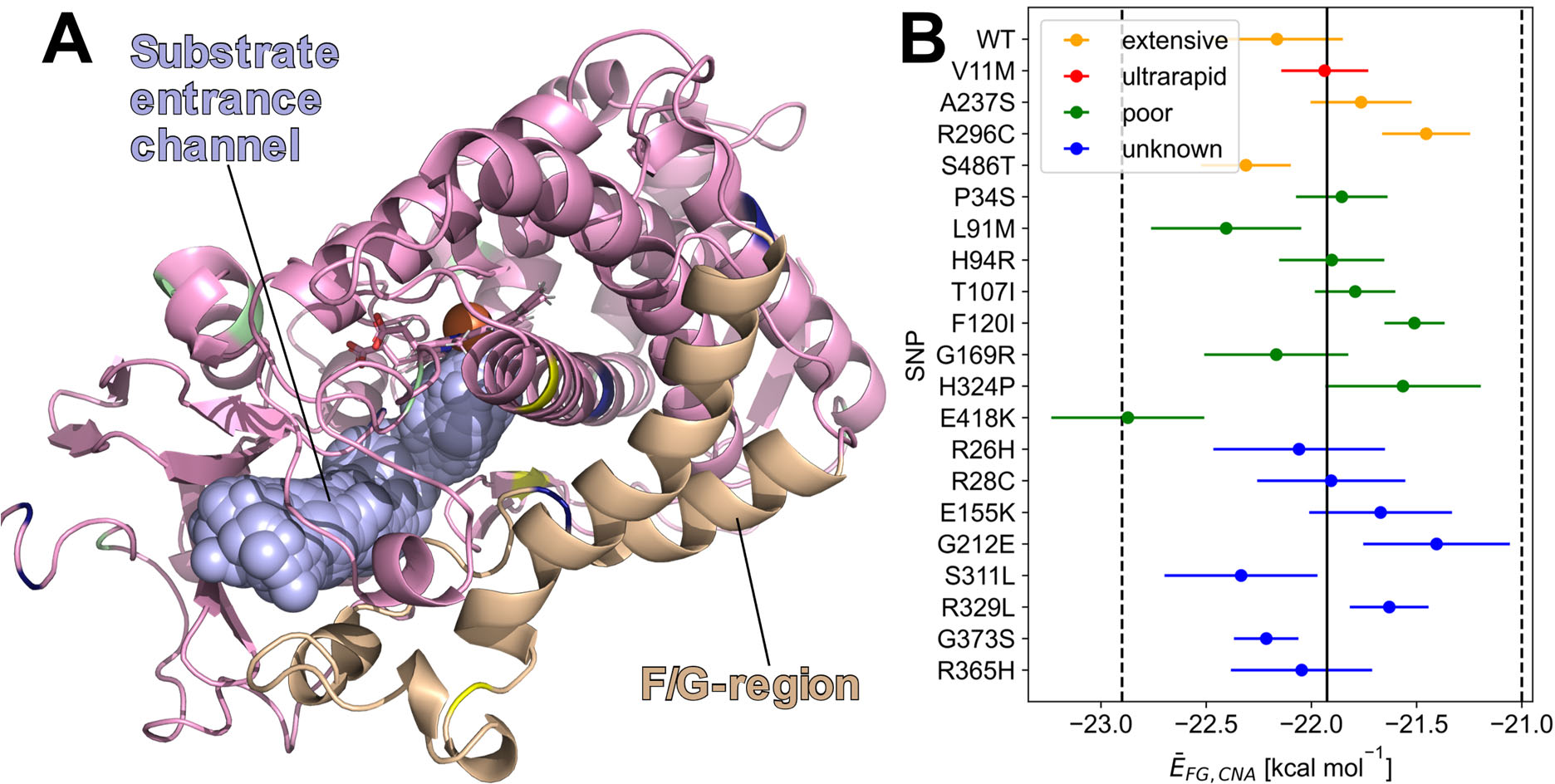
The structural stability of the F/G-region does not differ significantly between the different variants. **A:** The position of the substrate entrance channel (cyan) in CYP2D6 is next to the F/G region(beige). The rigidity of the F/G-region is correlated with substrate promiscuity in human CYP enzymes.^29^ Other parts of CYP2D6 are colored pink. Substitutions that lead to poor metabolism are colored in green, substitutions leading to an extensive metabolism are colored in yellow, V11M leading to an ultra-rapid metabolization is colored in red, and substitutions with unknown effects are colored in blue. **B:** To estimate the substrate promiscuity of the different isoforms, the structural stability of the F/G region (*Ē*_FG,CNA_) was calculated^29^. The continuous vertical line denotes the mean of *Ē_FG,_*_CNA_ of the CG; the error bars denote the SEM determined by error propagation along 10 independent simulations. The dashed vertical line denotes the limit for a significantly different value (*p* < 0.01, two-sided *t*-test). Since no substitution leads to a significant difference, the substrate promiscuity of the variants is likely similar to the wild type.

Here, we hypothesized that the overall reactivity of CYP2D6 can be decreased by substitutions that increase the structural stability of the F/G-region, as this would hamper substrate access due to a less mobile lid function. We, thus, computed *Ē*_FG,CNA_ for all variants with the same approach used before.^29^ However, differences between any of the variants and the WT were insignificant (Figure 6B). As a corollary of this finding, we expect that the substrate promiscuity of all investigated variants is similar to the wildtype.

### Decision tree to predict substitutions leading to poor metabolism

The above-analyzed aspects relevant to the CYP2D6-catalyzed reaction can be arranged in a hierarchy with associated criteria for when an aspect is significantly different from the CG. The resulting decision tree (Figure 7A) shall allow us to predict when a CYP2D6 substitution leads to poor metabolism.

**Figure 7:**
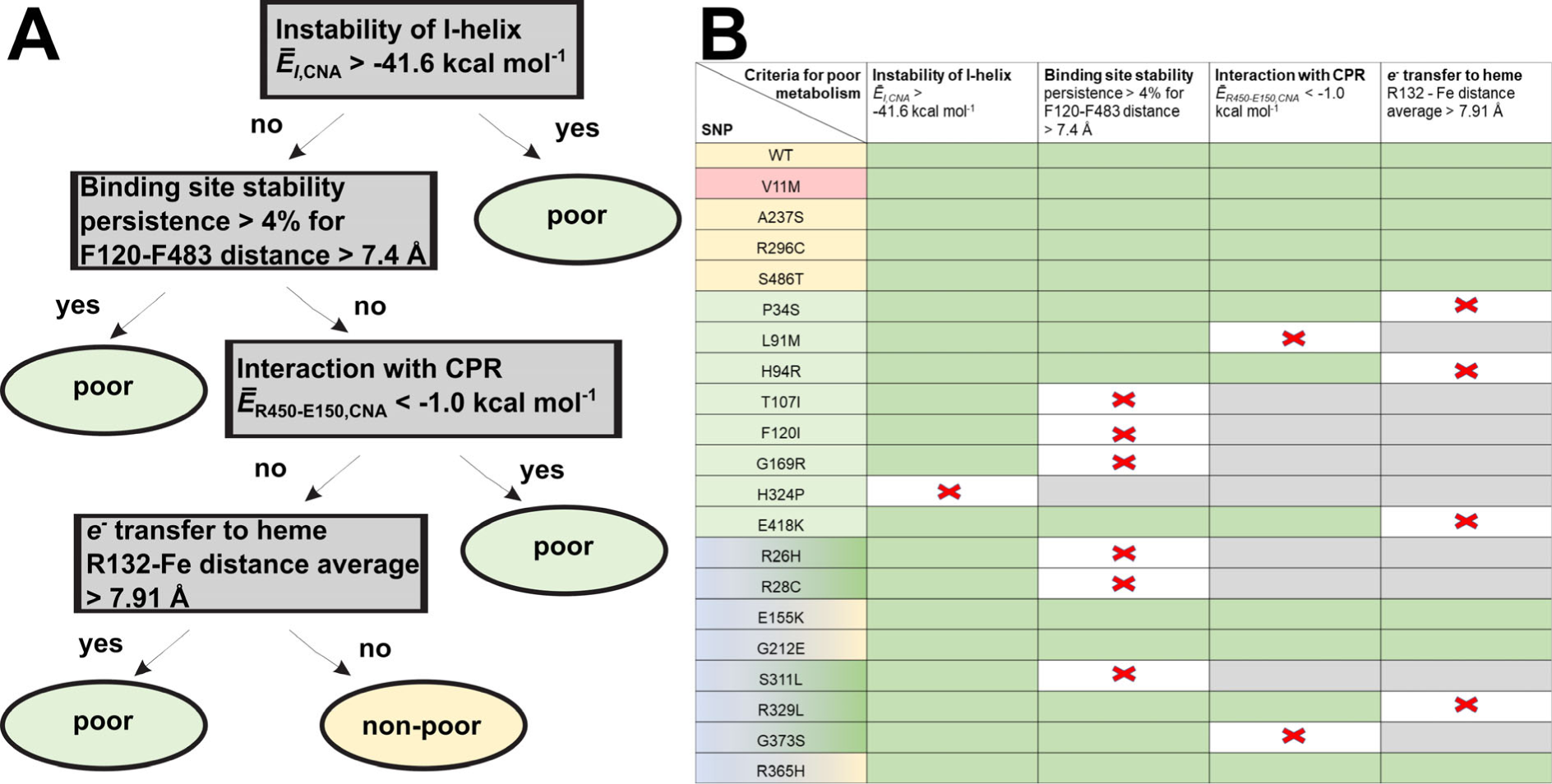
Decision tree for application of the derived criteria to the variants with unknown effect. **A:** Decision tree to distinguish poorly metabolizing variants from the CG. **B:** Aspects that lead to the prediction of a poor metabolism are marked with a red cross. Aspects that were not met are visualized with a green background, and aspects that were not checked due to the hierarchy of the decision tree are marked with a grey background. As a result, R26H, R28C, and S311L are predicted to lead to poor metabolism due to diminished binding site stability, whereas G373S is predicted to lead to poor metabolism due to diminished interaction with CPR. The first column is colored according to the (predicted) effect on the metabolism (poor metabolism: green; variants with no effect: yellow; V11M leading to an ultra-rapid metabolization: red; variants with unknown effects are colored with a color gradient from blue to the predicted effect on metabolism (green: poor; yellow: non-poor).

The importance of the I-helix as the central helix of the enzyme entails that if the enzyme is destabilized in this region, it loses its function.^71^ Thus, we use the structural stability of the I-helix as the most relevant aspect to determine if CYP2D6 activity is reduced. Compared to the CG of five variants with extensive or ultrarapid activity, an average chemical potential (eq. 2) in this region *Ē*_I,CNA_*^lim.^* = -41.57 kcal mol^-1^ is significantly different even when considering the maximum standard deviation of the nonCG (*p* < 0.01, two-sided t-test) (Figure 2B). Thus, variants with *E*_I,CNA_ > *E*_I,CNA_*^lim.^* are considered to lead to poor metabolism.

The substrate binding is a crucial step in the reaction process, which we consider the second most relevant aspect to determine if CYP2D6 activity is reduced. In the crystal structure of CYP2D6 bound to BACE1 (PDB ID: 4XRY), F120 and F483 stabilize the aromatic substrate (Figure 3A), and the distance between the ring systems of both residues is 7.4 Å. Assuming that this distance is relevant for substrate binding, we investigated the persistence with which this distance is above 7.4 Å during our MD trajectories. Compared to the CG, a persistence of 4% is significantly different (*p* < 0.01, two-sided t-test) (Figure 3B). Thus, variants with a persistence > 4% are considered to lead to poor metabolism.

The transfer of an electron from CPR is mandatory for the reaction cycle of CYP enzymes. Binding between both proteins is a prerequisite, which we consider the third most relevant aspect to determine if CYP2D6 activity is reduced. The unavailability of R450 of CPY2D6 as an interaction partner in the interface due to intramolecular salt bridge formation with E150 was analyzed in terms of the average chemical potential (eq. 2) of this interaction, *Ē*^R450^–^E150,CNA^. Compared to the CG, *Ē*_R450-E150,CNA_*^lim.^* = -1.0 kcal mol^-1^ is significantly different (*p* < 0.01, two-sided t-test) (Figure 4D). Thus, variants with *E*_R450-E150,CNA_ < *E*_R450-E150,CNA_*^lim.^* are considered to lead to poor metabolism.

We considered the transfer of an electron mediated by R132 as the last aspect to determine if CYP2D6 activity is reduced. As a criterion, the distance between iron and R132 was evaluated. Compared to the CG, a distance of 7.91 Å is significantly different (*p* < 0.01, two-sided t-test) (Figure 5C). Thus, variants with a distance > 7.91 Å are considered to lead to poor metabolism.

Following this decision tree, all variants of the CG and all variants that lead to known poor metabolism are correctly classified, with the first to fourth aspect leading to a decision in 1, 3, 1, and 3 cases for the latter group, respectively (Figure 7B). Application of this decision tree to variants with unknown effects leads to the predictions that variants R26H, R28C, S311L, R329L, and G373S result in poor metabolism.

### Possible molecular explanations for variants predicted to result in poor metabolism

#### Distortion of binding site integrity

R26H and R28C are part of the loop between the TM-helix and the A-helix and lead to a destabilization of the binding site as measured by the persistence with which the distance between F120 and F483 is above 7.4 Å during our MD trajectories. As residues 26 and 28 are at least 21.2 Å away from residues 120 and 483, this involves combined structural changes. F120 is pulled from the binding site due to a B’-helix (residues 105-109) movement (Figure S9A,B). The B’-helix moves due to a shift of the F’-helix. The shift is identified based on the constant distance between the B’- and F’-helices – measured as the distance between V223 (chosen as a reference point because V223 is the most central residue in the F’-helix) and Q108 (Figure S9C). The movement of the F’-helix is induced by a shift of the β-turn at W75. W75 is part of a β-turn between β1-1 strand and β1-2 strand, and the shift lets the F’-helix move toward the TM-helix (Figure S9D). The β-turn shifts due to a - in both variants -more frequently occurring hydrogen bond between the sidechain oxygen of Q27 and the indol nitrogen of W75 (Figure S9E,F).

S311L is part of the I-helix (Figure S10A) and leads to a destabilization of the binding site as measured by the persistence with which the distance between F120 and F483 is above 7.4 Å during our MD trajectories. F483 is moving away from the binding site because of a shift of the C-terminal loop, which contains F483. The shift is induced by W316 in the I-helix, which forms CH-π-stacking interactions with P487 in the C-terminal loop (Figure S10B,C). W316 is shifted by the increased spatial requirement of leucine compared to serine at position 311. The S311L substitution does not impact the I-helix itself, e.g., the distance between C_α_ 311 and C_α_ 316 does not change.

#### Reduced interaction with Cytochrome P450 reductase

G373S is part of the K-K’-loop, located between the K-helix and β1-4-strand. The K-K’-loop is close to the carboxy groups of heme (Figure S11A). The substitution is predicted to lead to a reduced interaction of the variant with CPR as indicated by *Ē*_R450-E150,CNA_ < -1.0 kcal mol^-^_1._

Due to a shift of E446 towards S373 in G373S, R450 is shifted closer to E150. The shift of E446 is induced by a shift of the K’-L-loop due to a decreased interaction between S437 and the heme carboxy group (Figure S11B). The distance between the S437 hydroxyl group, located in the K’-L-loop, and the heme carboxy group is significantly decreased by 0.3 Å compared to the CG (Figure S11C) (*p* < 0.01, two-sided *t*-test). The interaction between S437 and the heme carboxy group is decreased due to the frequently formed hydrogen bond between S373 in G373S, which cannot be formed in the GC.

#### Reduced electron transfer to heme mediated by R132

R329L is part of the J-helix and points to the protein surface. In the R329L variant, we observed a increase of *Ē*_J,CNA_ (eq. 2) of 0.6 kcal mol^-1^. Thus, the J-helix forms less stable rigid contacts to the surrounding structural elements. This leads to higher values of *Ē*_J-K’-loop,CNA_ and *Ē*_C,CNA_, which explains the higher mobility of R132 in the C-helix.

## Discussion

In this study, we intended to elucidate the effect of mutations found in naturally occurring alleles on CYP2D6 activity on an atomistic level. Our results demonstrate that SNPs correlated to changes in enzymatic activity led to changes in four crucial aspects of the CYP-catalyzed reaction, the stability of the main helix I-helix, substrate binding, binding of CPR, and electron transfer from CPR to heme iron. By contrast, no significant impacts on the substrate uptake channels due to the investigated SNPs were found.

We identified the four mechanisms by the most extensive MD simulations on CYP2D6 variants performed so far. Overall, we performed full atomistic MD simulations of membrane-bound CYP2D6 wildtype and variants of a cumulated time of 210 µs. This value exceeds previous studies^22, 72, 73^ by two orders of magnitude. The generated data allowed us to identify significant changes, even if they are small, among the different isoforms and versus the wildtype and to predict the behavior of unknown SNPs. For all four mechanisms, we identified parameters that can be used to distinguish between substitutions leading to a change in activity or not as well as are connected to the underlying molecular mechanisms. Also, the criteria match the current knowledge of the mechanism CYP enzymes undergo while oxidizing a substrate^12,13, 74^

To overcome the challenge of 165 known natural variants of CYP2D6, we investigated the 20 most frequent SNPs. This approach provided detailed insights into their molecular mechanisms but precludes to detect combinations of SNPs that lead to a larger structural change, e.g., CYP2D6*53, that combines F120I and A122S and does not show a change in activity^75^, even if F120I in CYP2D6*49 is associated with a decreased activity,^76^ and F120I was predicted as a mutation that decreases activity by HGMD^77^.

Interestingly, according to our analyses, the F/G-region stability and, by this, the substrate promiscuity is not changed by any of the investigated SNPs. This fits with the clinical practice because if a patient is identified as a poor metabolizer, metabolism will be poor for any CYP2D6 substrate.^78^ Thus, the drug regime will need to be modified once for all substrates and not for all substrates separately.

The derived mechanisms and the criteria associated with the mechanisms can be used to identify the effects of unknown SNPs and to scrutinize all known variants as to the molecualr origins that lead to the clinical effect. This can especially be useful for clinical effects with low incidence.

In summary, we identified four distinct mechanisms that lead to poor metabolism of CYP2D6 variants with the help of extensive MD simulations and Constraint Network Analysis. Our model shall allow us to predict if newly identified SNPs of CYP2D6 will lead to poor metabolism, which may be used for recommendations to modify drug regimes.

## Acknowledgments

We are grateful for computational support and infrastructure provided by the "Zentrum für Informations-und Medientechnologie" (ZIM) at the Heinrich Heine University Düsseldorf and the computing time provided by the John von Neumann Institute for Computing (NIC) on the supercomputer JUWELS at Jülich Supercomputing Centre (JSC) (user ID: HKF7, CYP450). The study was supported by Bundesministerium für Bildung und Forschung, Germany (BMBF) and the Department of Biotechnology, India (DBT) through project ReMetaDrug (funding number: 01DQ19002).

The FP7 WeNMR (project# 261572), H2020 West-Life (project# 675858), EOSC-hub (project# 777536), and the EGI-ACE (project# 101017567) European e-Infrastructure projects are acknowledged for the use of their web portals, which make use of the EGI infrastructure with the dedicated support of CESNET-MCC, INFN-PADOVA-STACK, INFN-LNL-2, NCG-INGRID-PT, TW-NCHC, CESGA, IFCA-LCG2, UA-BITP, SURFsara and NIKHEF, and the additional support of the national GRID Initiatives of Belgium, France, Italy, Germany, the Netherlands, Poland, Portugal, Spain, UK, Taiwan, and the US Open Science Grid.

## Author contributions

HG designed the study; DB performed computations; DB and HG analyzed results; DB and HG wrote the manuscript; PVB revised the manuscript; HG and PVB secured funding.

## Data and software availability

For molecular simulations, the AMBER18 package of molecular simulation codes was used. AMBER18 is available from here: http://ambermd.org/.

The CNA software is available under academic licenses from http://cpclab.uni-duesseldorf.de/index.php/Software. The CNA web server is accessible at http://cpclab.uni-duesseldorf.de/cna/.

## References

1. Rendic, S.; Guengerich, F. P., Survey of Human Oxidoreductases and Cytochrome P450 Enzymes Involved in the Metabolism of Xenobiotic and Natural Chemicals. Chem. Res. Toxicol. 2015, 28, 38–42.

2. Mahgoub, A.; Dring, L. G.; Idle, J. R.; Lancaster, R.; Smith, R. L., POLYMORPHIC HYDROXYLATION OF DEBRISOQUINE IN MAN. The Lancet 1977, 310, 584–586.

3. Kimura, S.; Umeno, M.; Skoda, R. C.; Meyer, U. A.; Gonzalez, F. J., The human debrisoquine 4-hydroxylase (CYP2D) locus: sequence and identification of the polymorphic CYP2D6 gene, a related gene, and a pseudogene. Am. J. Hum. Genet. 1989, 45, 889–904.

4. Nebert, D. W.; Russell, D. W., Clinical importance of the cytochromes P450. Lancet 2002, 360, 1155–1162.

5. Dahl, M. L.; Johansson, I.; Bertilsson, L.; Ingelman-Sundberg, M.; Sjöqvist, F., Ultrarapid hydroxylation of debrisoquine in a Swedish population. Analysis of the molecular genetic basis. J. Pharmacol. Exp. Ther. 1995, 274, 516–20.

6. Løvlie, R.; Daly, A. K.; Molven, A.; Idle, J. R.; Steen, V. M., Ultrarapid metabolizers of debrisoquine: Characterization and PCR-based detection of alleles with duplication of the CYP2D6 gene. FEBS Lett. 1996, 392, 30–34.

7. Johansson, I.; Lundqvist, E.; Bertilsson, L.; Dahl, M. L.; Sjöqvist, F.; Ingelman-Sundberg, M., Inherited amplification of an active gene in the cytochrome P450 CYP2D locus as a cause of ultrarapid metabolism of debrisoquine. Proceedings of the National Academy of Sciences 1993, 90, 11825–11829.

8. Chiba, K.; Kato, M.; Ito, T.; Suwa, T.; Sugiyama, Y., Inter-individual variability of *in vivo* CYP2D6 activity in different genotypes. Drug Metabolism and Pharmacokinetics 2012, advpub, 1201160353–1201160353.

9. Yu, A.; Kneller, B. M.; Rettie, A. E.; Haining, R. L., Expression, Purification, Biochemical Characterization, and Comparative Function of Human Cytochrome P450 2D6.1, 2D6.2, 2D6.10, and 2D6.17 Allelic Isoforms. J. Pharmacol. Exp. Ther. 2002, 303, 1291.

10. Guengerich, F. P. Human Cytochrome P450 Enzymes. In *Cytochrome P450*: Structure, Mechanism, and Biochemistry, Ortiz de Montellano, P. R., Ed.; Springer International Publishing: Cham, 2015, pp 523–785.

11. Black, S. D., Membrane topology of the mammalian P450 cytochromes. FASEB J. 1992, 6, 680–685.

12. Cojocaru, V.; Winn, P. J.; Wade, R. C., The ins and outs of cytochrome P450s. *Biochim. Biophys. Acta*, Gen. Subj. 2007, 1770, 390–401.

13. Schlichting, I.; Berendzen, J.; Chu, K.; Stock Ann, M.; Maves Shelley, A.; Benson David, E.; Sweet Robert, M.; Ringe, D.; Petsko Gregory, A.; Sligar Stephen, G., The Catalytic Pathway of Cytochrome P450cam at Atomic Resolution. Science 2000, 287, 1615–1622.

14. Masters, B. S. S.; Okita, R. T., The history, properties, and function of NADPH-cytochrome P-450 reductase. Pharmacol. Ther. 1980, 9, 227–244.

15. Hamdane, D.; Xia, C.; Im, S.-C.; Zhang, H.; Kim, J.-J. P.; Waskell, L., Structure and Function of an NADPH-Cytochrome P450 Oxidoreductase in an Open Conformation Capable of Reducing Cytochrome P450 *. J. Biol. Chem. 2009, 284, 11374–11384.

16. Prade, E.; Mahajan, M.; Im, S.-C.; Zhang, M.; Gentry, K. A.; Anantharamaiah, G. M.; Waskell, L.; Ramamoorthy, A., A Minimal Functional Complex of Cytochrome P450 and FBD of Cytochrome P450 Reductase in Nanodiscs. Angew. Chem. Int. Ed. 2018, 57, 8458–8462.

17. Ortiz de Montellano, P. R. Substrate Oxidation by Cytochrome P450 Enzymes. In Cytochrome P450: Structure, Mechanism, and Biochemistry, Ortiz de Montellano, P. R., Ed.; Springer International Publishing: Cham, 2015, pp 111–176.

18. Xia, C.; Hamdane, D.; Shen, A. L.; Choi, V.; Kasper, C. B.; Pearl, N. M.; Zhang, H.; Im, S.-C.; Waskell, L.; Kim, J.-J. P., Conformational Changes of NADPH-Cytochrome P450 Oxidoreductase Are Essential for Catalysis and Cofactor Binding *. J. Biol. Chem. 2011, 286, 16246–16260.

19. Waskell, L.; Kim, J.-J. P. Electron Transfer Partners of Cytochrome P450. In Cytochrome P450: Structure, Mechanism, and Biochemistry, Ortiz de Montellano, P. R., Ed.; Springer International Publishing: Cham, 2015, pp 33–68.

20. Dai, D.-P.; Geng, P.-W.; Wang, S.-H.; Cai, J.; Hu, L.-M.; Nie, J.-J.; Hu, J.-H.; Hu, G.-X.; Cai, J.-P., In vitro functional assessment of 22 newly identified CYP2D6 allelic variants in the Chinese population. Basic Clin Pharmacol Toxicol 2015, 117, 39–43.

21. de Groot, M. J.; Wakenhut, F.; Whitlock, G.; Hyland, R., Understanding CYP2D6 interactions. Drug Discov. Today 2009, 14, 964–972.

22. Huff, H. C.; Vasan, A.; Roy, P.; Kaul, A.; Tajkhorshid, E.; Das, A., Differential Interactions of Selected Phytocannabinoids with Human CYP2D6 Polymorphisms. Biochemistry 2021, 60, 2749–2760.

23. Zanger, U. M.; Fischer, J.; Raimundo, S.; Stüven, T.; Evert, B. O.; Schwab, M.; Eichelbaum, M., Comprehensive analysis of the genetic factors determining expression and function of hepatic CYP2D6. Pharmacogenetics and Genomics 2001, 11.

24. Flanagan, J. U.; MarÉChal, J.-D.; Ward, R.; Kemp, C. A.; McLaughlin, L. A.; Sutcliffe, M. J.; Roberts, G. C. K.; Paine, M. J. I.; Wolf, C. R., Phe120 contributes to the regiospecificity of cytochrome P450 2D6: mutation leads to the formation of a novel dextromethorphan metabolite. Biochem. J. 2004, 380, 353–360.

25. Keizers, P. H. J.; Lussenburg, B. M. A.; de Graaf, C.; Mentink, L. M.; Vermeulen, N. P. E.; Commandeur, J. N. M., Influence of phenylalanine 120 on cytochrome P450 2D6 catalytic selectivity and regiospecificity: crucial role in 7-methoxy-4-(aminomethyl)-coumarin metabolism. Biochem. Pharmacol. 2004, 68, 2263–2271.

26. Lussenburg, B. M. A.; Keizers, P. H. J.; de Graaf, C.; Hidestrand, M.; Ingelman-Sundberg, M.; Vermeulen, N. P. E.; Commandeur, J. N. M., The role of phenylalanine 483 in cytochrome P450 2D6 is strongly substrate dependent. Biochem. Pharmacol. 2005, 70, 1253–1261.

27. Gaedigk, A.; Ingelman-Sundberg, M.; Miller, N. A.; Leeder, J. S.; Whirl-Carrillo, M.; Klein, T. E.; the PharmVar Steering, C., The Pharmacogene Variation (PharmVar) Consortium: Incorporation of the Human Cytochrome P450 (CYP) Allele Nomenclature Database. Clinical Pharmacology & Therapeutics 2018, 103, 399–401.

28. Mulnaes, D.; Porta, N.; Clemens, R.; Apanasenko, I.; Reiners, J.; Gremer, L.; Neudecker, P.; Smits, S. H. J.; Gohlke, H., TopModel: Template-Based Protein Structure Prediction at Low Sequence Identity Using Top-Down Consensus and Deep Neural Networks. J. Chem. Theory Comput. 2020, 16, 1953–1967.

29. Becker, D.; Bharatam, P. V.; Gohlke, H., F/G Region Rigidity is Inversely Correlated to Substrate Promiscuity of Human CYP Isoforms Involved in Metabolism. J. Chem. Inf. Model. 2021, 61, 4023–4030.

30. Jumper, J.; Evans, R.; Pritzel, A.; Green, T.; Figurnov, M.; Ronneberger, O.; Tunyasuvunakool, K.; Bates, R.; Žídek, A.; Potapenko, A.; Bridgland, A.; Meyer, C.; Kohl, S. A. A.; Ballard, A. J.; Cowie, A.; Romera-Paredes, B.; Nikolov, S.; Jain, R.; Adler, J.; Back, T.; Petersen, S.; Reiman, D.; Clancy, E.; Zielinski, M.; Steinegger, M.; Pacholska, M.; Berghammer, T.; Bodenstein, S.; Silver, D.; Vinyals, O.; Senior, A. W.; Kavukcuoglu, K.; Kohli, P.; Hassabis, D., Highly accurate protein structure prediction with AlphaFold. Nature 2021, 596, 583–589.

31. System, T. P. M. G. The PyMOL Molecular Graphics System, Version 2.0; Schrödinger, LLC.

32. Søndergaard, C. R.; Olsson, M. H. M.; Rostkowski, M.; Jensen, J. H., Improved Treatment of Ligands and Coupling Effects in Empirical Calculation and Rationalization of pKa Values. J. Chem. Theory Comput. 2011, 7, 2284–2295.

33. Doerr, S.; Harvey, M. J.; Noé, F.; De Fabritiis, G., HTMD: High-Throughput Molecular Dynamics for Molecular Discovery. J. Chem. Theory Comput. 2016, 12, 1845–1852.

34. Schott-Verdugo, S.; Gohlke, H., PACKMOL-Memgen: A Simple-To-Use, Generalized Workflow for Membrane-Protein-Lipid-Bilayer System Building. J. Chem. Inf. Model. 2019, 59, 2522–2528.

35. Yeagle, P., The membranes of cells. Third edition. ed.; Elsevier/AP: Amsterdam ; Boston, 2016; p xii, 439 pages.

36. Nugent, T.; Jones, D. T., Membrane protein orientation and refinement using a knowledge-based statistical potential. BMC Bioinformatics 2013, 14, 276.

37. Salomon-Ferrer, R.; Götz, A. W.; Poole, D.; Le Grand, S.; Walker, R. C., Routine Microsecond Molecular Dynamics Simulations with AMBER on GPUs. 2. Explicit Solvent Particle Mesh Ewald. J. Chem. Theory Comput. 2013, 9, 3878–3888.

38. Case, D. A.; Ben-Shalom, I. Y.; Brozell, S. R.; Cerutti, D. S.; Cheatham, I. T.E.; Cruzeiro, V. W. D.; Darden, T. A.; Duke, R. E.; Ghoreishi, D.; Giambasu, G.; Giese, T.; Gilson, M. K.; Gohlke, H.; Goetz, A. W.; Greene, D.; Harris, R.; Homeyer, N.; Huang, Y.; Izadi, S.; Kovalenko, A.; Krasny, R.; Kurtzman, T.; Lee, T. S.; LeGrand, S.; Li, P.; Lin, C.; Liu, J.; Luchko, T.; Luo, R.; Man, V.; Mermelstein, D. J.; Merz, K. M.; Miao, Y.; Monard, G.; Nguyen, C.; Nguyen, H.; Onufriev, A.; Pan, F.; Qi, R.; Roe, D. R.; Roitberg, A.; Sagui, C.; Schott-Verdugo, S.; Shen, J.; Simmerling, C. L.; Smith, J.; Swails, J.; Walker, R. C.; Wang, J.; Wei, H.; Wilson, L.; Wolf, R. M.; Wu, X.; Xiao, L.; Xiong, Y.; York, D. M.; Kollman, P. A. *AMBER* 2019, University of California, San Francisco, 2019.

39. Maier, J. A.; Martinez, C.; Kasavajhala, K.; Wickstrom, L.; Hauser, K. E.; Simmerling, C., ff14SB: Improving the Accuracy of Protein Side Chain and Backbone Parameters from ff99SB. J. Chem. Theory Comput. 2015, 11, 3696–3713.

40. Dickson, C. J.; Madej, B. D.; Skjevik, A. A.; Betz, R. M.; Teigen, K.; Gould, I. R.; Walker, R. C., Lipid14: The Amber Lipid Force Field. J. Chem. Theory Comput. 2014, 10, 865–879.

41. Izadi, S.; Anandakrishnan, R.; Onufriev, A. V., Building Water Models: A Different Approach. J. Phys. Chem. Lett. 2014, 5, 3863–3871.

42. Sengupta, A.; Li, Z.; Song, L. F.; Li, P.; Merz, K. M., Jr., Parameterization of Monovalent Ions for the OPC3, OPC, TIP3P-FB, and TIP4P-FB Water Models. J. Chem. Inf. Model. 2021, 61, 869–880.

43. Shahrokh, K.; Orendt, A.; Yost, G. S.; Cheatham, T. E., 3rd, Quantum mechanically derived AMBER-compatible heme parameters for various states of the cytochrome P450 catalytic cycle. J. Comput. Chem. 2012, 33, 119–133.

44. Ryckaert, J.-P.; Ciccotti, G.; Berendsen, H. J. C., Numerical integration of the cartesian equations of motion of a system with constraints: molecular dynamics of n-alkanes. J. Comput. Phys. 1977, 23, 327–341.

45. Schott-Verdugo, S.; Muller, L.; Classen, E.; Gohlke, H.; Groth, G., Structural Model of the ETR1 Ethylene Receptor Transmembrane Sensor Domain. Sci Rep 2019, 9, 8869.

46. Quigley, D.; Probert, M. I. J., Langevin dynamics in constant pressure extended systems. J. Chem. Phys. 2004, 120, 11432–11441.

47. Berendsen, H. J. C.; Postma, J. P. M.; van Gunsteren, W. F.; DiNola, A.; Haak, J. R., Molecular dynamics with coupling to an external bath. J. Chem. Phys. 1984, 81, 3684–3690.

48. Roe, D. R.; Cheatham, T. E., PTRAJ and CPPTRAJ: Software for Processing and Analysis of Molecular Dynamics Trajectory Data. J. Chem. Theory Comput. 2013, 9, 3084–3095.

49. Case, D. A.; Aktulga, H. M.; Belfon, K.; Cerutti, D. S.; Cisneros, G. A.; Cruzeiro, V. W. D.; Forouzesh, N.; Giese, T. J.; Götz, A. W.; Gohlke, H.; Izadi, S.; Kasavajhala, K.; Kaymak, M. C.; King, E.; Kurtzman, T.; Lee, T.-S.; Li, P.; Liu, J.; Luchko, T.; Luo, R.; Manathunga, M.; Machado, M. R.; Nguyen, H. M.; O’Hearn, K. A.; Onufriev, A. V.; Pan, F.; Pantano, S.; Qi, R.; Rahnamoun, A.; Risheh, A.; Schott-Verdugo, S.; Shajan, A.; Swails, J.; Wang, J.; Wei, H.; Wu, X.; Wu, Y.; Zhang, S.; Zhao, S.; Zhu, Q.; Cheatham, T. E., III; Roe, D. R.; Roitberg, A.; Simmerling, C.; York, D. M.; Nagan, M. C.; Merz, K. M., Jr., AmberTools. J. Chem. Inf. Model. 2023.

50. Hermans, S. M. A.; Pfleger, C.; Nutschel, C.; Hanke, C. A.; Gohlke, H., Rigidity theory for biomolecules: concepts, software, and applications. Wiley Interdiscip. Rev.: Comput. Mol. Sci. 2017, 7, e1311.

51. Jacobs, D. J.; Rader, A. J.; Kuhn, L. A.; Thorpe, M. F., Protein flexibility predictions using graph theory. Proteins 2001, 44, 150–165.

52. Dahiyat, B. I.; Benjamin Gordon, D.; Mayo, S. L., Automated design of the surface positions of protein helices. Protein Sci. 1997, 6, 1333–1337.

53. Radestock, S.; Gohlke, H., Exploiting the Link between Protein Rigidity and Thermostability for Data-Driven Protein Engineering. Eng. Life Sci. 2008, 8, 507–522.

54. Pfleger, C.; Rathi, P. C.; Klein, D. L.; Radestock, S.; Gohlke, H., Constraint Network Analysis (CNA): A Python Software Package for Efficiently Linking Biomacromolecular Structure, Flexibility, (Thermo-)Stability, and Function. J. Chem. Inf. Model. 2013, 53, 1007–1015.

55. Jacobs, D. J.; Thorpe, M. F., Generic Rigidity Percolation: The Pebble Game. Phys. Rev. Lett. 1995, 75, 4051–4054.

56. Pfleger, C.; Radestock, S.; Schmidt, E.; Gohlke, H., Global and local indices for characterizing biomolecular flexibility and rigidity. J. Comput. Chem. 2013, 34, 220–233.

57. Pfleger, C.; Minges, A.; Boehm, M.; McClendon, C. L.; Torella, R.; Gohlke, H., Ensemble- and Rigidity Theory-Based Perturbation Approach To Analyze Dynamic Allostery. J. Chem. Theory Comput. 2017, 13, 6343–6357.

58. van Zundert, G. C. P.; Rodrigues, J. P. G. L. M.; Trellet, M.; Schmitz, C.; Kastritis, P. L.; Karaca, E.; Melquiond, A. S. J.; van Dijk, M.; de Vries, S. J.; Bonvin, A. M. J. J., The HADDOCK2.2 Web Server: User-Friendly Integrative Modeling of Biomolecular Complexes. J. Mol. Biol. 2016, 428, 720–725.

59. Honorato, R. V.; Koukos, P. I.; Jiménez-García, B.; Tsaregorodtsev, A.; Verlato, M.; Giachetti, A.; Rosato, A.; Bonvin, A. M. J. J., Structural Biology in the Clouds: The WeNMR-EOSC Ecosystem. Frontiers in Molecular Biosciences 2021, 8.

60. UniProt, C., UniProt: a worldwide hub of protein knowledge. Nucleic Acids Res. 2019, 47, D506–D515.

61. Sherry, S. T.; Ward, M. H.; Kholodov, M.; Baker, J.; Phan, L.; Smigielski, E. M.; Sirotkin, K., dbSNP: the NCBI database of genetic variation. Nucleic Acids Res. 2001, 29, 308–11.

62. Landrum, M. J.; Chitipiralla, S.; Brown, G. R.; Chen, C.; Gu, B.; Hart, J.; Hoffman, D.; Jang, W.; Kaur, K.; Liu, C.; Lyoshin, V.; Maddipatla, Z.; Maiti, R.; Mitchell, J.; O’Leary, N.; Riley, G. R.; Shi, W.; Zhou, G.; Schneider, V.; Maglott, D.; Holmes, J. B.; Kattman, B. L., ClinVar: improvements to accessing data. Nucleic Acids Res. 2020, 48, D835–D844.

63. HGMD (human gene mutation database). In Encyclopedia of Genetics, Genomics, Proteomics and Informatics, Rédei, G. P., Ed.; Springer Netherlands: Dordrecht, 2008, pp 873–873.

64. Virtanen, P.; Gommers, R.; Oliphant, T. E.; Haberland, M.; Reddy, T.; Cournapeau, D.; Burovski, E.; Peterson, P.; Weckesser, W.; Bright, J.; van der Walt, S. J.; Brett, M.; Wilson, J.; Millman, K. J.; Mayorov, N.; Nelson, A. R. J.; Jones, E.; Kern, R.; Larson, E.; Carey, C. J.; Polat, *İ*.; Feng, Y.; Moore, E. W.; VanderPlas, J.; Laxalde, D.; Perktold, J.; Cimrman, R.; Henriksen, I.; Quintero, E. A.; Harris, C. R.; Archibald, A. M.; Ribeiro, A. H.; Pedregosa, F.; van Mulbregt, P.; Vijaykumar, A.; Bardelli, A. P.; Rothberg, A.; Hilboll, A.; Kloeckner, A.; Scopatz, A.; Lee, A.; Rokem, A.; Woods, C. N.; Fulton, C.; Masson, C.; Häggström, C.; Fitzgerald, C.; Nicholson, D. A.; Hagen, D. R.; Pasechnik, D. V.; Olivetti, E.; Martin, E.; Wieser, E.; Silva, F.; Lenders, F.; Wilhelm, F.; Young, G.; Price, G. A.; Ingold, G.-L.; Allen, G. E.; Lee, G. R.; Audren, H.; Probst, I.; Dietrich, J. P.; Silterra, J.; Webber, J. T.; Slavič, J.; Nothman, J.; Buchner, J.; Kulick, J.; Schönberger, J. L.; de Miranda Cardoso, J. V.; Reimer, J.; Harrington, J.; Rodríguez, J. L. C.; Nunez-Iglesias, J.; Kuczynski, J.; Tritz, K.; Thoma, M.; Newville, M.; Kümmerer, M.; Bolingbroke, M.; Tartre, M.; Pak, M.; Smith, N. J.; Nowaczyk, N.; Shebanov, N.; Pavlyk, O.; Brodtkorb, P. A.; Lee, P.; McGibbon, R. T.; Feldbauer, R.; Lewis, S.; Tygier, S.; Sievert, S.; Vigna, S.; Peterson, S.; More, S.; Pudlik, T.; Oshima, T.; Pingel, T. J.; Robitaille, T. P.; Spura, T.; Jones, T. R.; Cera, T.; Leslie, T.; Zito, T.; Krauss, T.; Upadhyay, U.; Halchenko, Y. O.; Vázquez-Baeza, Y.; SciPy, C., SciPy 1.0: fundamental algorithms for scientific computing in Python. Nat. Methods 2020, 17, 261–272.

65. Morris, A. L.; MacArthur, M. W.; Hutchinson, E. G.; Thornton, J. M., Stereochemical quality of protein structure coordinates. Proteins 1992, 12, 345–364.

66. Ramesh, M.; Bharatam, P. V., Importance of hydrophobic parameters in identifying appropriate pose of CYP substrates in cytochromes. Eur. J. Med. Chem. 2014, 71, 15–23.

67. Roughley, S. D.; Jordan, A. M., The Medicinal Chemist’s Toolbox: An Analysis of Reactions Used in the Pursuit of Drug Candidates. J. Med. Chem. 2011, 54, 3451–3479.

68. Im, S.-C.; Waskell, L., The interaction of microsomal cytochrome P450 2B4 with its redox partners, cytochrome P450 reductase and cytochrome b5. Arch. Biochem. Biophys. 2011, 507, 144–153.

69. Bridges, A.; Gruenke, L.; Chang, Y.-T.; Vakser, I. A.; Loew, G.; Waskell, L., Identification of the Binding Site on Cytochrome P450 2B4 for Cytochrome <em>b</em> 5 and Cytochrome P450 Reductase *. J. Biol. Chem. 1998, 273, 17036–17049.

70. Kumar, S.; Nussinov, R., Close-Range Electrostatic Interactions in Proteins. ChemBioChem 2002, 3, 604–617.

71. Evert, B.; Griese, E.-U.; Eichelbaum, M., A missense mutation in exon 6 of the CYP2D6 gene leading to a histidine 324 to proline exchange is associated with the poor metabolizer phenotype of sparteine. Naunyn-Schmiedeberg’s Arch. Pharmacol. 1994, 350, 434–439.

72. Fischer, A.; Don, C. G.; Smieško, M., Molecular Dynamics Simulations Reveal Structural Differences among Allelic Variants of Membrane-Anchored Cytochrome P450 2D6. J. Chem. Inf. Model. 2018, 58, 1962–1975.

73. Dorne, J. L. C. M.; Cirlini, M.; Louisse, J.; Pedroni, L.; Galaverna, G.; Dellafiora, L., A Computational Understanding of Inter-Individual Variability in CYP2D6 Activity to Investigate the Impact of Missense Mutations on Ochratoxin A Metabolism. 2022, 14, 207.

74. Shaik, S.; Dubey, K. D., The catalytic cycle of cytochrome P450: a fascinating choreography. Trends in Chemistry 2021, 3, 1027–1044.

75. Ebisawa, A.; Hiratsuka, M.; Sakuyama, K.; Konno, Y.; Sasaki, T.; Mizugaki, M., Two novel single nucleotide polymorphisms (SNPs) of the CYP2D6 gene in Japanese individuals. Drug Metab Pharmacokinet 2005, 20, 294–9.

76. Soyama, A.; Kubo, T.; Miyajima, A.; Saito, Y.; Shiseki, K.; Komamura, K.; Ueno, K.; Kamakura, S.; Kitakaze, M.; Tomoike, H.; Ozawa, S.; Sawada, J., Novel nonsynonymous single nucleotide polymorphisms in the CYP2D6 gene. Drug Metab Pharmacokinet 2004, 19, 313–9.

77. Masayuki, M.; Hiroshi, Y.; Kazuma, K.; Shunsuke, I.; Jyunji, S.; Kazuko, N.; Akiko, S.; Shogo, O.; Jun-Ichi, S.; Eiji, K.; Moritoshi, K.; Tetsuya, K., Two Novel <em>CYP2D6*10</em> Haplotypes As Possible Causes of a Poor Metabolic Phenotype in Japanese. Drug Metab. Dispos. 2009, 37, 699.

78. Crews, K. R.; Monte, A. A.; Huddart, R.; Caudle, K. E.; Kharasch, E. D.; Gaedigk, A.; Dunnenberger, H. M.; Leeder, J. S.; Callaghan, J. T.; Samer, C. F.; Klein, T. E.; Haidar, C. E.; Van Driest, S. L.; Ruano, G.; Sangkuhl, K.; Cavallari, L. H.; Müller, D. J.; Prows, C. A.; Nagy, M.; Somogyi, A. A.; Skaar, T. C., Clinical Pharmacogenetics Implementation Consortium Guideline for CYP2D6, OPRM1, and COMT Genotypes and Select Opioid Therapy. Clinical Pharmacology & Therapeutics 2021, 110, 888–896.

