## Supporting Information for "Molecular mechanisms underlying single nucleotide polymorphism-induced reactivity decrease in CYP2D6"

### Supporting Methods

#### Statistical Analysis and Data Analysis of Computational Results

The standard error of the mean (SEM) for the computed $\bar{E}_{region,\mathrm{CNA}}$ values was calculated by considering each trajectory as an independent sample. Data of CNA analyses were visualized by visualCNA.[1]

The SEM was propagated by eq. S1:

$$SEM = \frac{1}{10}\sqrt{\sum_{i=1}^{10} SEM_i^{\,2}} \qquad \textbf{(eq. S1)}$$

All calculations were performed with NumPy.[2] Curve fitting was performed with the SciPy module stats.[3]

#### Linear Interaction Energy Analysis

In the analysis of the G169R variant, the linear interaction energy (LIE) was calculated by the *lie* function implemented in pytraj.[4] We[5] and others[6, 7] showed that LIE is an efficient method to obtain good affinity predictions. The electrostatic and van der Waals interactions of R169 were calculated to the surrounding residues S168 and P496. The cutoff for interactions was set to 12.0 Å. The ligand mask was set to the atoms of the guanidino group, and the surrounding mask was set to S168 and P496.

### Supplemental Figures

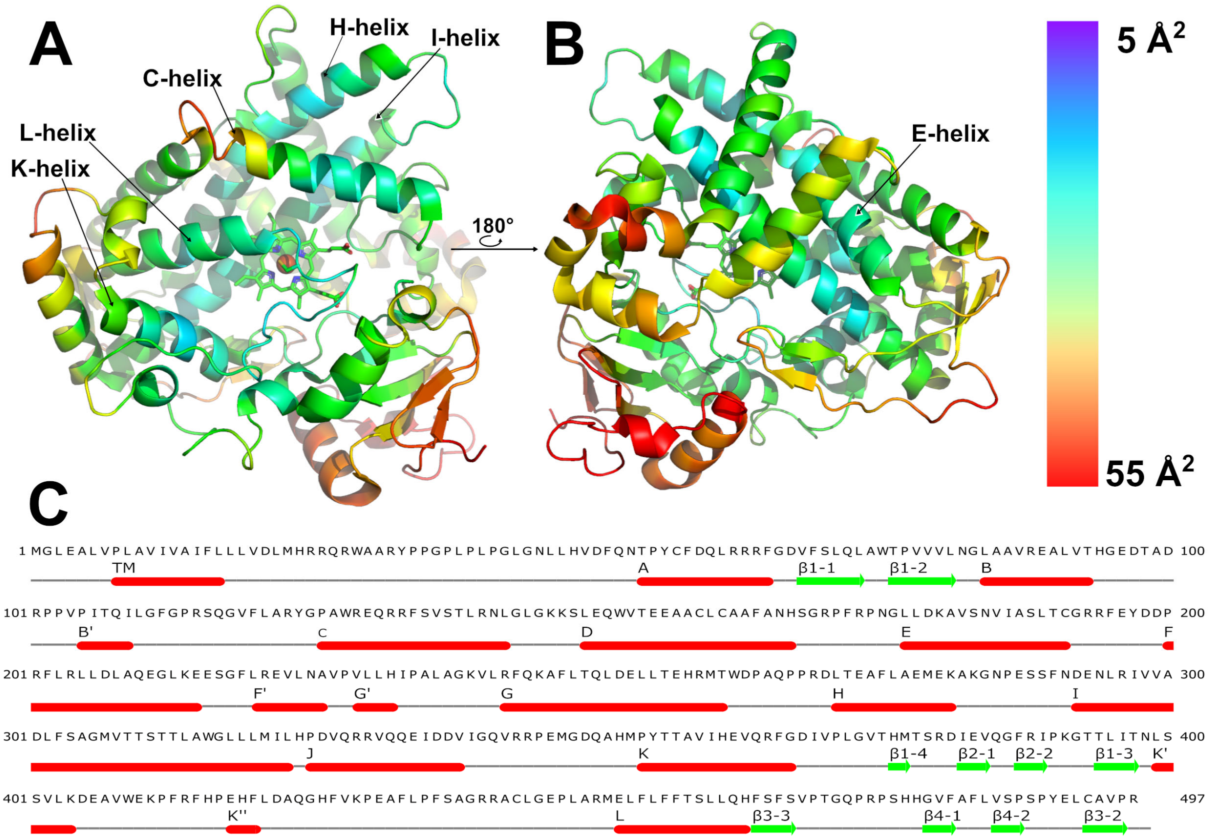


**Figure S1:** Crystal structure (PDB ID: 3TBG) of the globular part of CYP2D6 colored by B-factor (red: most mobile, blue: least mobile). **A:** The helices E, H, I, L, and K are the least mobile helices in CYP2D6. The heme ring system is shown as sticks, while the heme iron is shown as a sphere. **B:** Structure of CYP2D6 rotated by 180°. The color bar on the right denotes the B-factor. **C:** Sequence of CYP2D6 with marked secondary structure elements and the nomenclature.

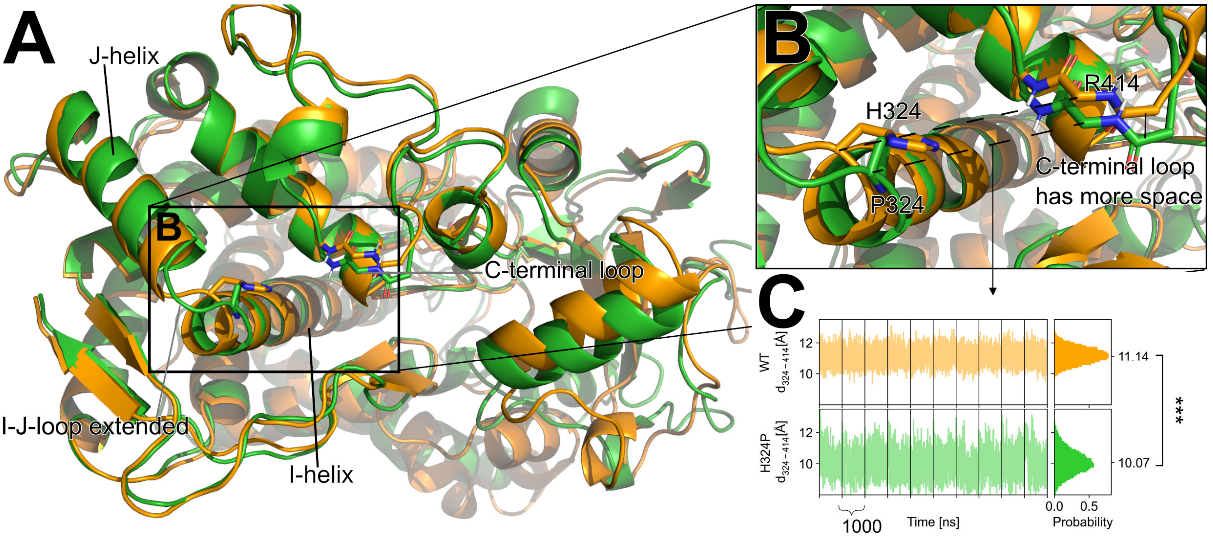


**Figure S2:** Effect of the H324P mutation on CYP2D6. **A:** The mutation in the variant H324P (lime-green) is located in the loop between I-helix and J-helix. Compared to the wild type (orange), the I-J-loop is extended by one residue. **B:** In variant H324P, the C-terminal loop (marked with an arrow) is shifted towards the I-helix. Sidechains at position 324 are shown as sticks, and the backbone of R414 is shown as sticks. Dashed lines indicate distances measured in panel C. **C:** The $C_\alpha$ distance between residues 324 and 414 is significantly decreased for the variant H324P compared to wild type. Left: time series of the distance, values per trajectory are separated by black vertical lines; right: aggregate probability distributions. Statistical analysis was performed using the two-sided t-test (* $p < 0.01$; ** $p < 0.001$; *** $p < 0.0001$; n.s.: $p >$ 0.01).

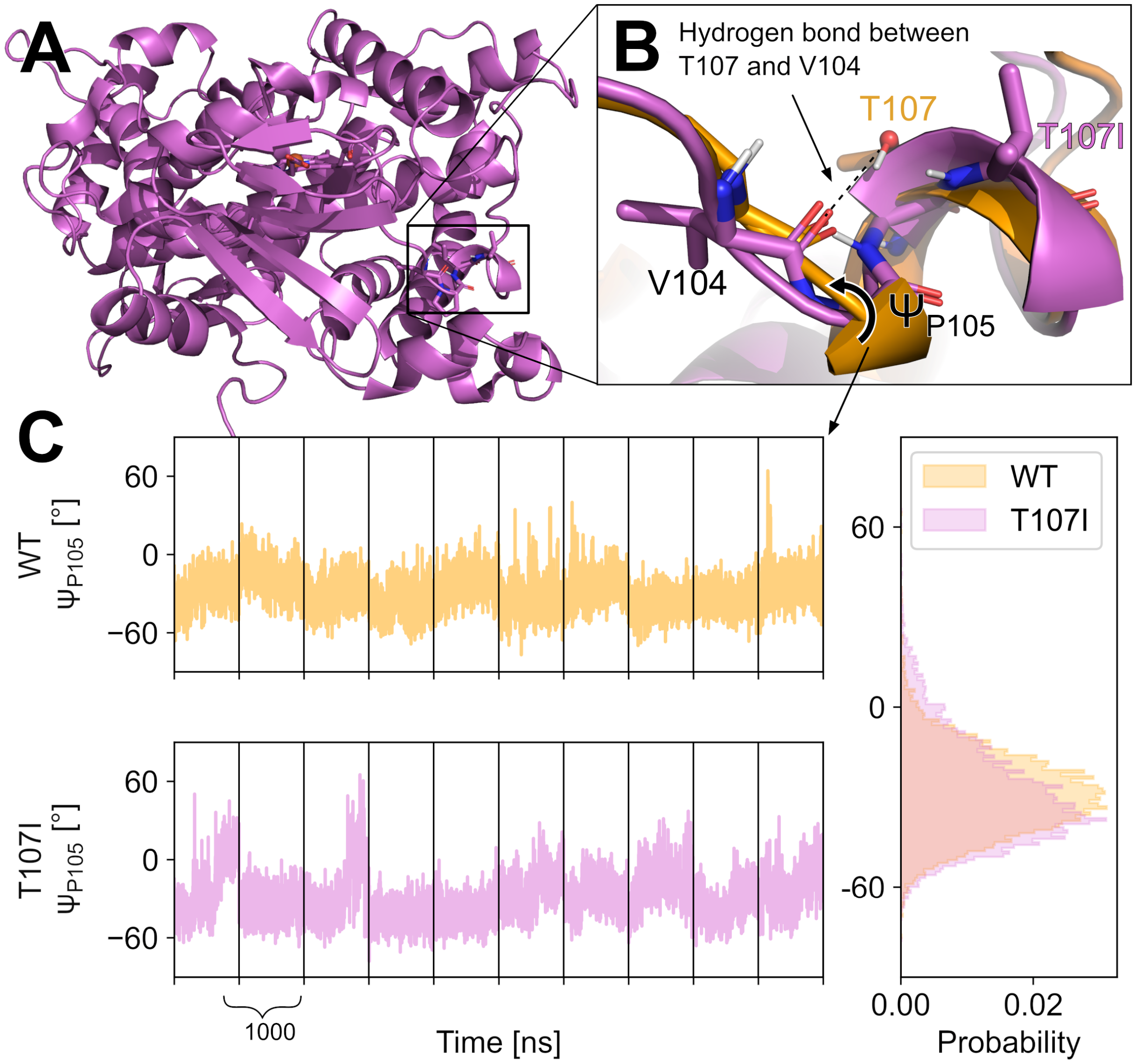


**Figure S3:** Effect of the T107I mutation on CYP2D6. **A:** The mutation in the variant T107I (orchid) is located in the C'-helix. The heme ring system is shown as sticks, while the heme iron is shown as a sphere. **B:** The missing interaction between V104 and T107 in the T107I variant destabilizes the C'-helix. The backbone of positions 104-107 is shown as sticks, as is the sidechain of position 107. The Ψ torsion at P105 is indicated with an arrow. **C:** The Ψ torsion has significantly more often positive values ($p < 0.01$, two-sided *t*-test) in T107I compared to the wild type. Left: time series of the Ψ torsion, values per trajectory are separated by black vertical lines; right: aggregate probability distributions.

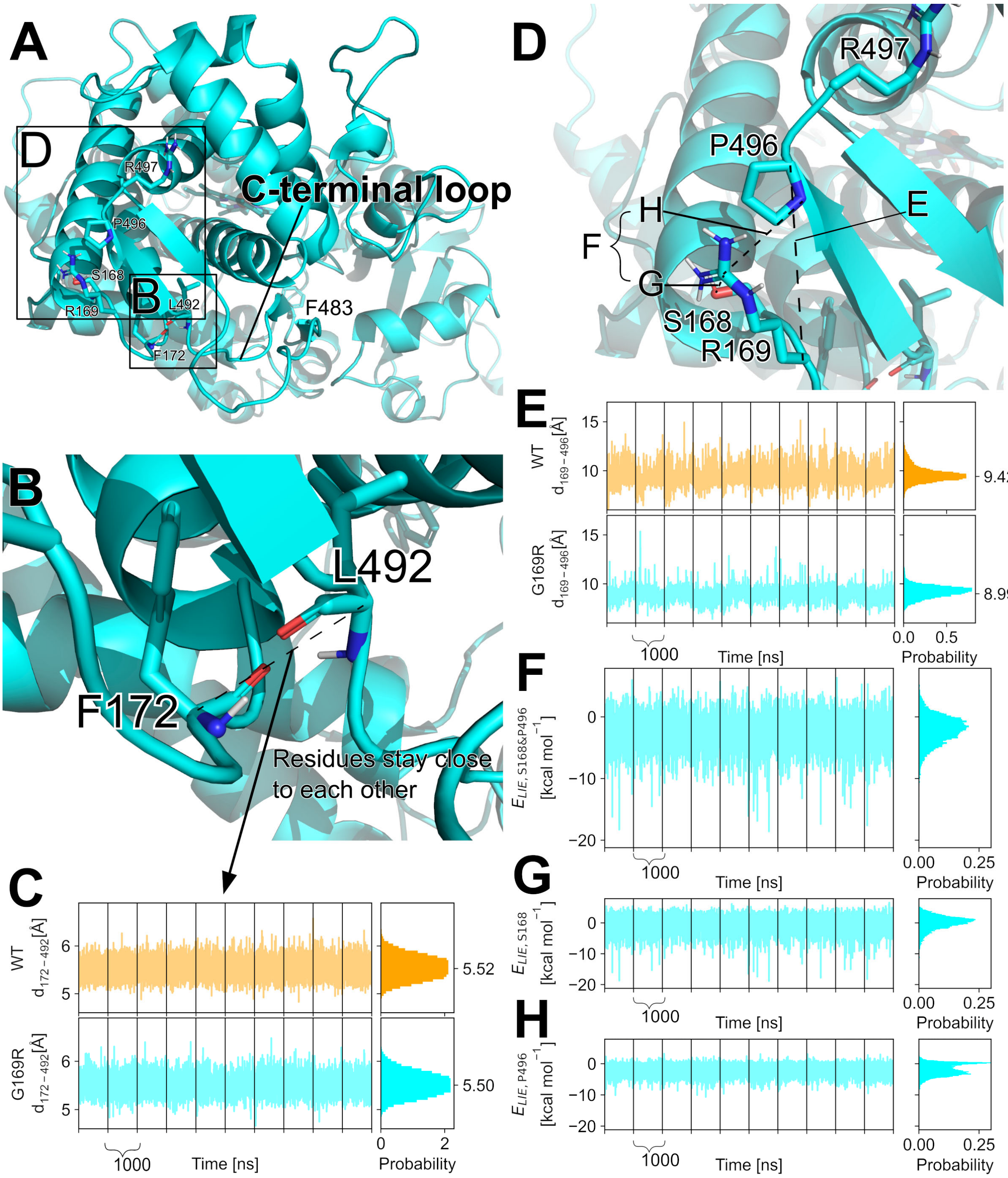


**Figure S4:** Effect of the G169R mutation on CYP2D6. **A:** The mutation in the variant G169R (cyan) is located in the D-E-loop. The heme ring system is shown as sticks, while the heme iron is shown as a sphere. **B:** F172 and L492 form backbone hydrogen atoms between the D-E-loop and the C-terminal loop. Amide hydrogens of both residues point to the carbonyl oxygens of the opposite residues. The backbone of both residues is shown as sticks. A dashed line indicates the distance measured in panel C. **C:** The $C_\alpha$ distance between F172 and L492 is highly constant and similar in the wild type (orange) and the variant G169R. Left: time series of the distance, values per trajectory are separated by black vertical lines; right: aggregate probability distributions. **D:** R169 in the variant G169R interacts with S168 and P496.

Interactions evaluated in panels F-H are indicated with dashed lines and the measured distance between $C_\alpha$ 169 and $C_\alpha$ 496 (panel E). **E:** The distance between $C_\alpha$ 169 and $C_\alpha$ 496 is significantly decreased in the variant G169R compared to wild type ($p < 0.01$, two-sided *t*-test). Left: time series of the distance, values per trajectory are separated by black vertical lines; right: aggregate probability distributions. **F-H:** Electrostatic and van der Waals energies computed according to the Linear Interaction Energy approach between the R169 guanidino group and S168 and P496 (F), R169 guanidino group and S168 (G), and R169 guanidino group and P496 (H) in the G169R variant. Left: time series of the electrostatic and van der Waals energies, values per trajectory are separated by black vertical lines; right: aggregate probability distributions. (F) In most frames, R169 interacts favorably with S168 *and* P496. In comparison to panels G and H, both residues interact with R169 in the investigated G169R variant. (G) In most frames, R169 shows interactions with S168. (H) In most frames, R169 shows interactions with P496.

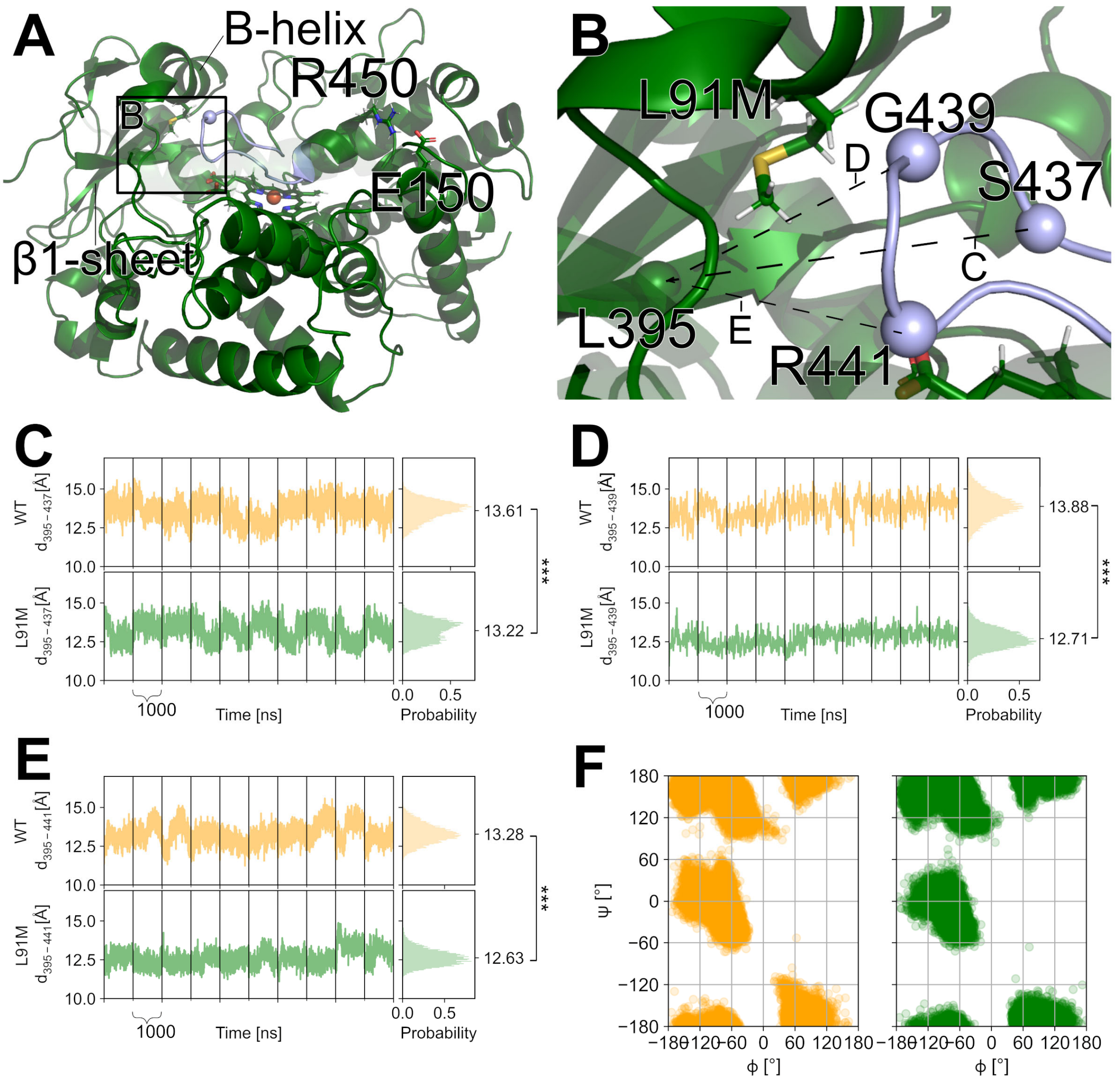


**Figure S5:** Effect of the L91M mutation on CYP2D6. **A:** The mutation L91M (green) is located on the B-helix. The heme ring is shown as sticks, while the heme iron and important $C_\alpha$ atoms are shown as spheres. The C-helix is shown with high transparency for visualization purposes. **B:** The missing second $C_\gamma$ in the variant leads to a closer distance between the K-L-loop (purple) and the β1 strand. $C_\alpha$ of L395 and G439 are shown as spheres. A dashed line indicates the measured distance. **C:** The distance between $C_\alpha$ atoms of L395 and S437 is significantly reduced in the L91M variant compared to wild type ($p < 0.01$, two-sided *t*-test). **D:** The distance between $C_\alpha$ atoms of L395 and G439 is significantly reduced in the L91M variant compared to wild type ($p < 0.01$, two-sided *t*-test). **E:** The distance between $C_\alpha$ atoms of L395 and R441 is significantly reduced in the L91M variant compared to wild type ($p < 0.01$, two-sided *t*-test). C-E: Left: time series of the distance, black vertical lines separate values per trajectory; right: aggregate probability distributions. **F:** Left: Ramachandran plot of the K-L-loop (residues 436-444) of the wild type. Right: Ramachandran plot of the K-L-loop (residues 436-444) of the L91M. Each data point represents one residue in one snapshot over 10 x 1 µs of simulation time. No major conformational change is visible between wild type and variant. Statistical analysis was performed using the two-sided t-test (* $p < 0.01$; ** $p < 0.001$; *** $p < 0.0001$; n.s.: $p > 0.01$).

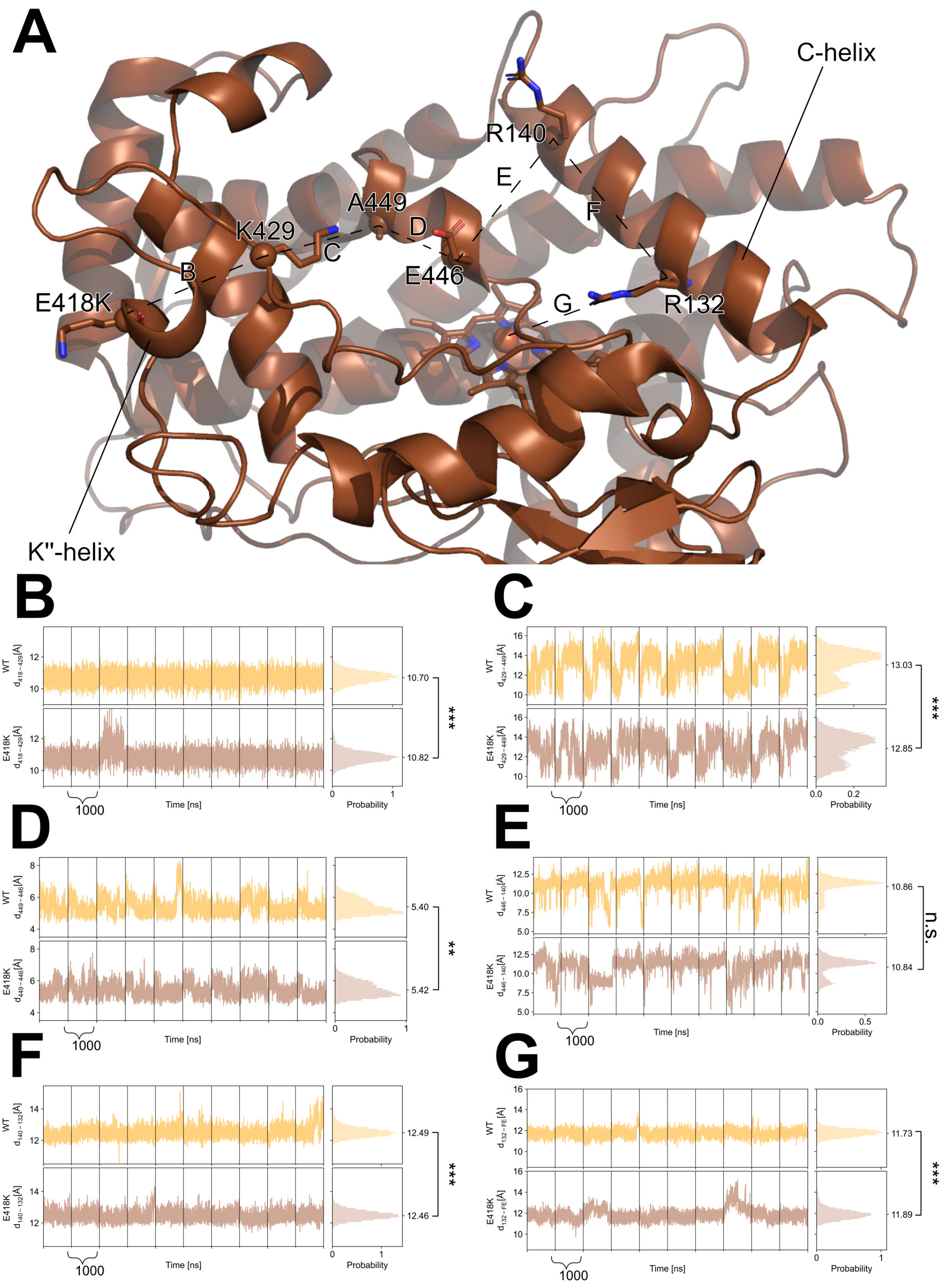


**Figure S6:** Effect of the E418K mutation on CYP2D6. **A:** The mutation in E418K is located on the K''-helix, while R132 is part of the C-helix. The $C_\alpha$ atoms of residues between a distance measured are shown as spheres. Dashed lines indicate measured distances. The heme ring system is shown as sticks, while the heme iron is shown as a sphere. **B:** The distance between

$C_\alpha$ of residue E418K and $C_\alpha$ of residue K429 is significantly increased in E418K ($p < 0.01$, two-sided *t*-test). **C:** The distance between $C_\alpha$ of residue K429 and $C_\alpha$ of residue A449 is significantly decreased in E418K ($p < 0.01$, two-sided *t*-test). **D:** The distance between $C_\alpha$ of residue E446 and $C_\alpha$ of residue A449 differs on average only by 0.02 Å. **E:** The distance between $C_\alpha$ of residue R140 and $C_\alpha$ of residue E446 does not differ significantly between WT and E418K. **F:** The distance between $C_\alpha$ of residue R132 and $C_\alpha$ of residue R140 differs on average only by 0.03 Å. **G:** The distance between $C_\alpha$ of residue R132 and iron in heme is significantly increased in E418K ($p < 0.01$, two-sided *t*-test). B-G: Left: time series of the distance, black vertical lines separate values per trajectory; right: aggregate probability distributions. Statistical analysis was performed using the two-sided t-test (* $p < 0.01$; ** $p < 0.001$; *** $p < 0.0001$; n.s.: $p > 0.01$).

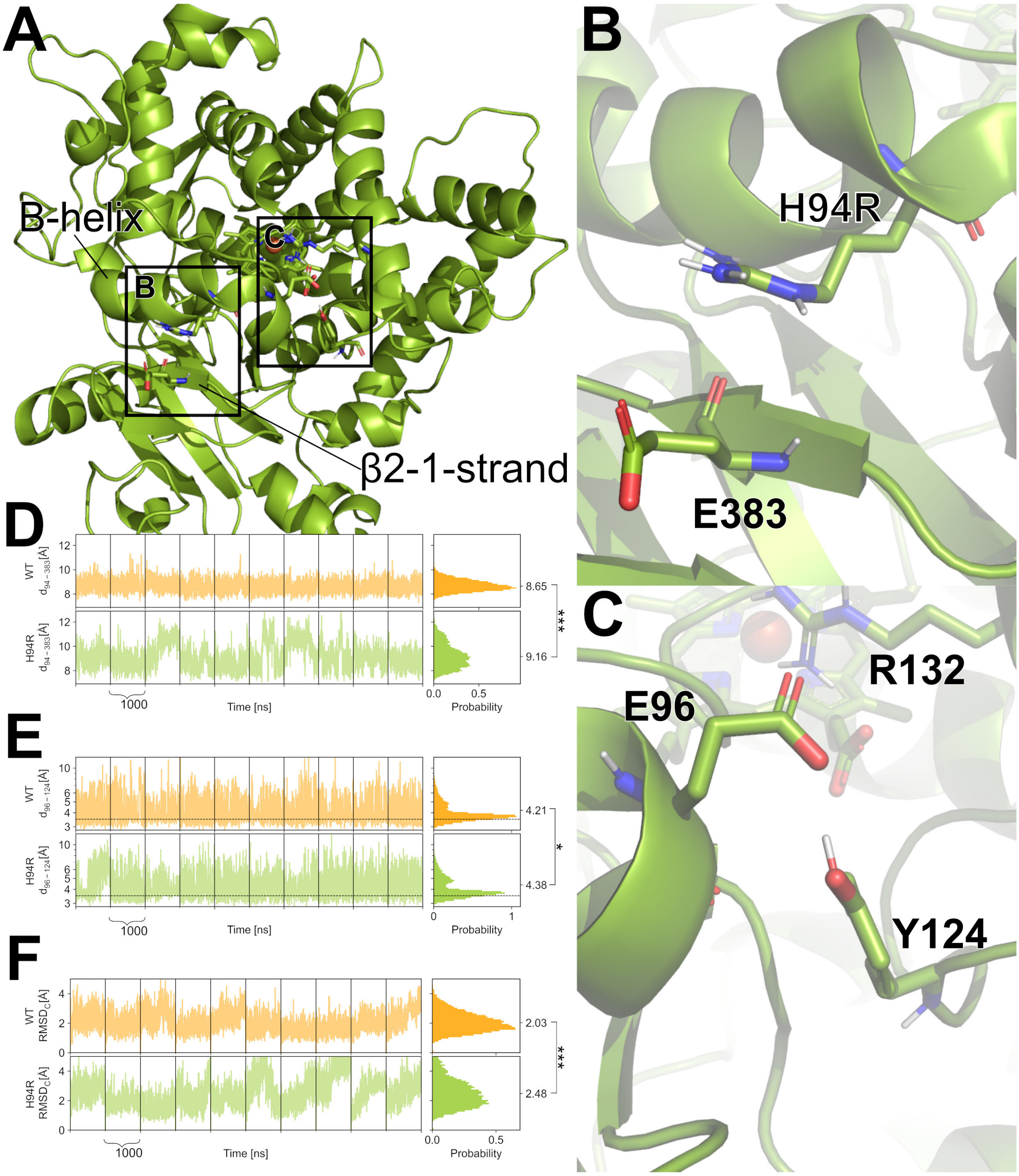


**Figure S7:** Effect of the H94R mutation on CYP2D6. **A:** The mutation in H94R is located on the B-helix. The heme ring and important residues are shown as sticks, while the heme iron is shown as a sphere. **B:** Due to the larger space requirement of R94, the distance between the Cα atom of R94 and the Cα atom of E383 is increased. E383 is part of the β2-1 strand. **C:** The charge-assisted hydrogen bond between E96 and Y124 is more often formed in WT than in H94R due to a shift of the B-helix. **D:** Distance between the Cα atom of residue H94/R94 and the Cα atom of residue E383 is significantly increased ($p < 0.01$, two-sided *t*-test). **E:** The hydrogen bond between E96 and Y124 is significantly less often formed in H94R than in the wild type ($p < 0.01$, two-sided *t*-test). The dashed line at 3.5 Å indicates the formation of a hydrogen bond. **F:** The $C_\alpha$-RMSD (residues 126-143) of the C-helix after fitting on the heme group is significantly increased. D-F: Left: time series of the distance or the $C_\alpha$-RMSD, values per trajectory are separated by black vertical lines; right: aggregate probability distributions. Statistical analysis was performed using the two-sided t-test (* $p < 0.01$;** $p < 0.001$; *** $p < 0.0001$; n.s.: $p > 0.01$).

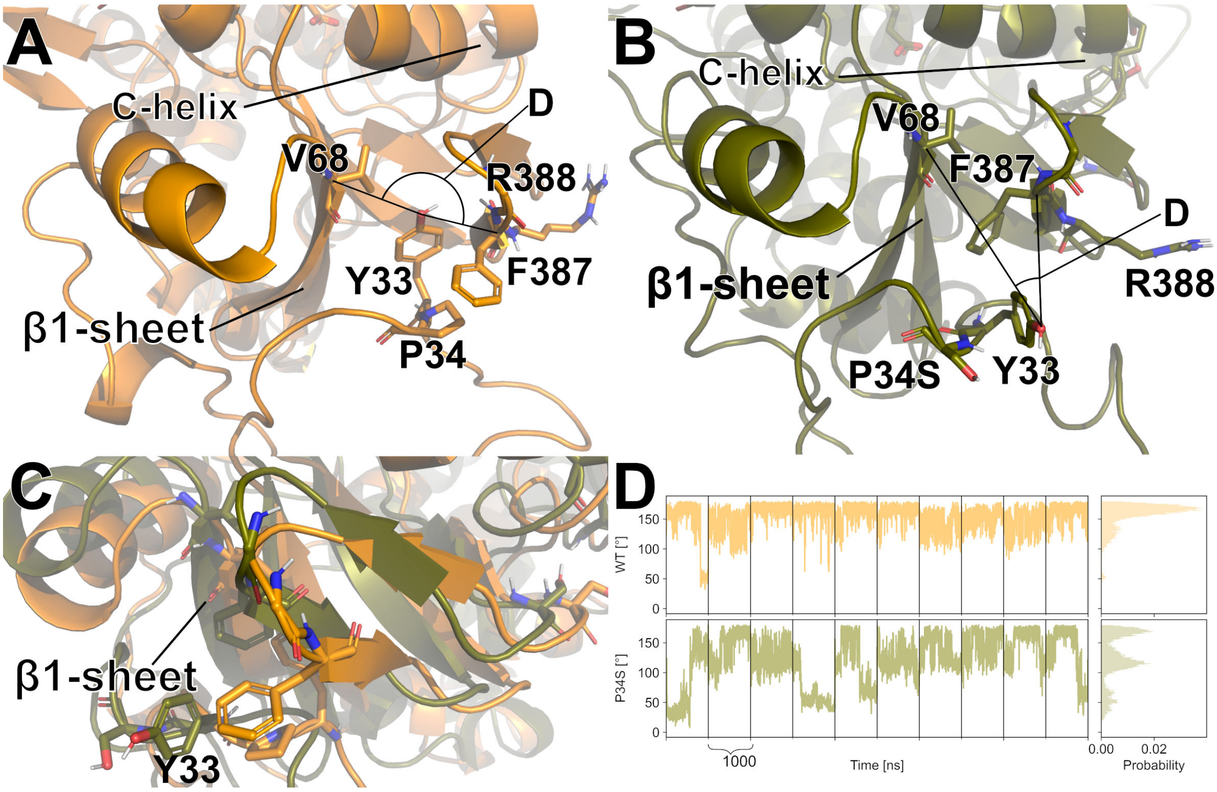


**Figure S8:** Effect of the P34S mutation on CYP2D6. **A:** P34 in CYP2D6 wild type stabilizes the position of Y33 by rigidifying the loop Y33 belongs to between V68 and F387 (see also panel D). Important residues are shown as sticks. Black lines indicate the angle measured in panel D. **B:** P34S in the CYP2D6 variant does not stabilize the position of Y33 between V68 and F387 (see also panel D). Y33 is shifted away from the β1 strand, and F387 is moving towards V69. Important residues are shown as sticks. Black lines indicate the angle measured in panel D. **C:** Overlay of CYP2D6 wild type (orange) and the P34S variant (olive) reveals the drastic change in the β1 strand conformation upon the Y33 shift. The relative orientation of the β1 strand leads to changes in the interaction with the C-helix. **D:** Y33 flips out of its position more often in P34S than in the wild type. Left: time series of the angle ($C_\alpha$ of V68, oxygen of the hydroxy group of Y33, $C_\alpha$ of F387), values per trajectory are separated by black vertical lines; right: aggregate probability distributions.

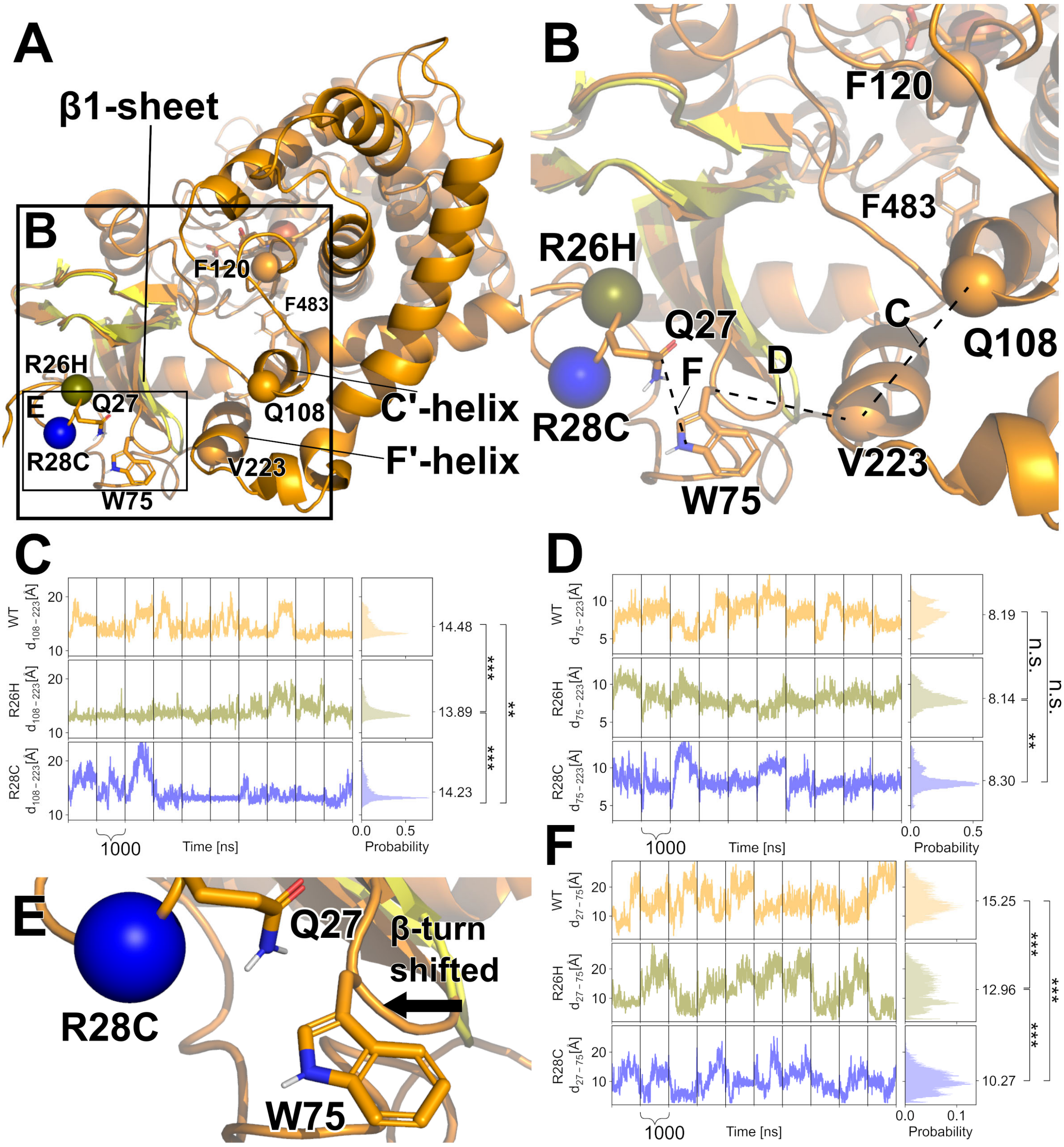


**Figure S9:** Effect of the R26H and R28C mutations on CYP2D6. **A:** The mutations R26H (olive) and R28C (blue) are located in the loop between the TM-helix and β1-sheet. The heme ring system is shown as sticks, while the heme iron is shown as a sphere. **B:** Mutations in the loop between the TM-helix and strand β1 influence the F'-helix and C'-helix. Q27 and W75 are shown as sticks; the $C_\alpha$ atoms of residues between a distance was measured are shown as spheres. Dashed lines indicate measured distances. **C:** The distance between $C_\alpha$ of residue Q108 and $C_\alpha$ of residue V223 is unaffected by either variant. **D:** The distance between $C_\alpha$ of residue W75 and $C_\alpha$ of residue V223 is significantly decreased in R26H and R28C ($p < 0.01$, two-sided *t*-test). **E:** Q27 interacts with W75 and leads to a shift of the β turn. **F:** The distance between $C_\alpha$ of residue Q27 and the aromatic nitrogen of W75 is significantly decreased in R26H and R28C

($p < 0.01$, two-sided *t*-test). C, D, F: Left: time series of the distance, values per trajectory are separated by black vertical lines; right: aggregate probability distributions. Statistical analysis was performed using the two-sided t-test (* $p < 0.01$;** $p < 0.001$; *** $p < 0.0001$; n.s.: $p > 0.01$).

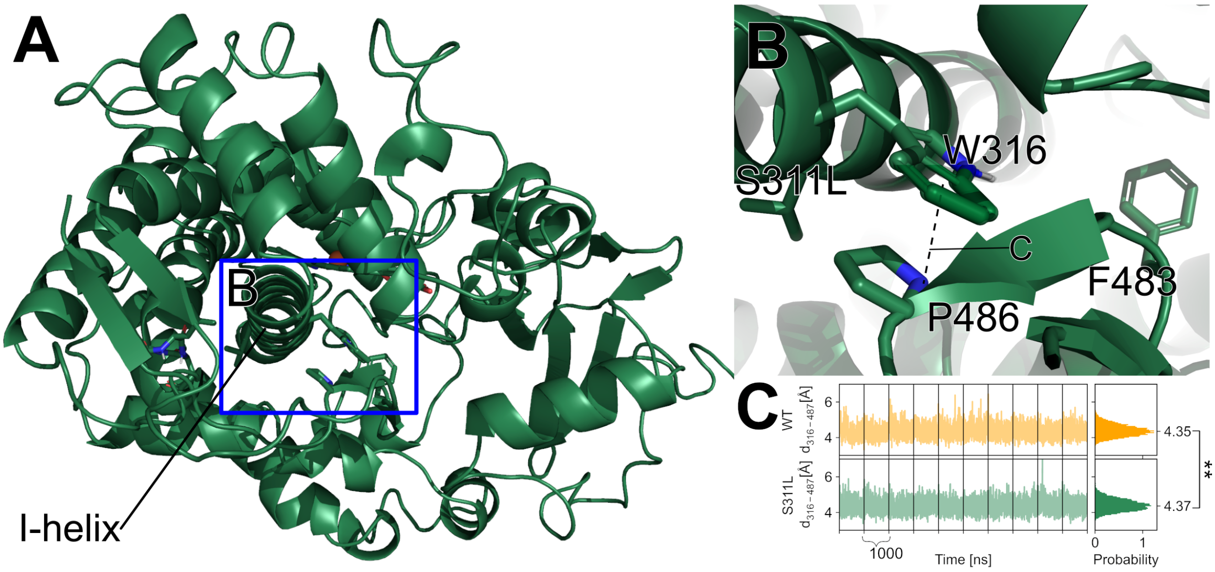


**Figure S10:** Effect of the S311L mutation on CYP2D6. **A:** The mutation S311L (sea green) is located in the I-helix. The heme ring system is shown as sticks, while the heme iron is shown as a sphere. **B:** W316 and P486 have a "stacked-like" arrangement.[8] The close contact pushes the C-terminal loop to the β1-sheet in the S311L variant due to space requirements. Residues S311L, W316, F483, and P486 are shown as sticks. Dashed lines indicate the measured distance. **C:** The distance between $C_\alpha$ of residue W316 and $C_\alpha$ of residue P486 differs on average only by 0.02 Å.. Left: time series of the distance, values per trajectory are separated by black vertical lines; right: aggregate probability distributions. Statistical analysis was performed using the two-sided t-test (* $p < 0.01$;** $p < 0.001$; *** $p < 0.0001$; n.s.: $p > 0.01$).

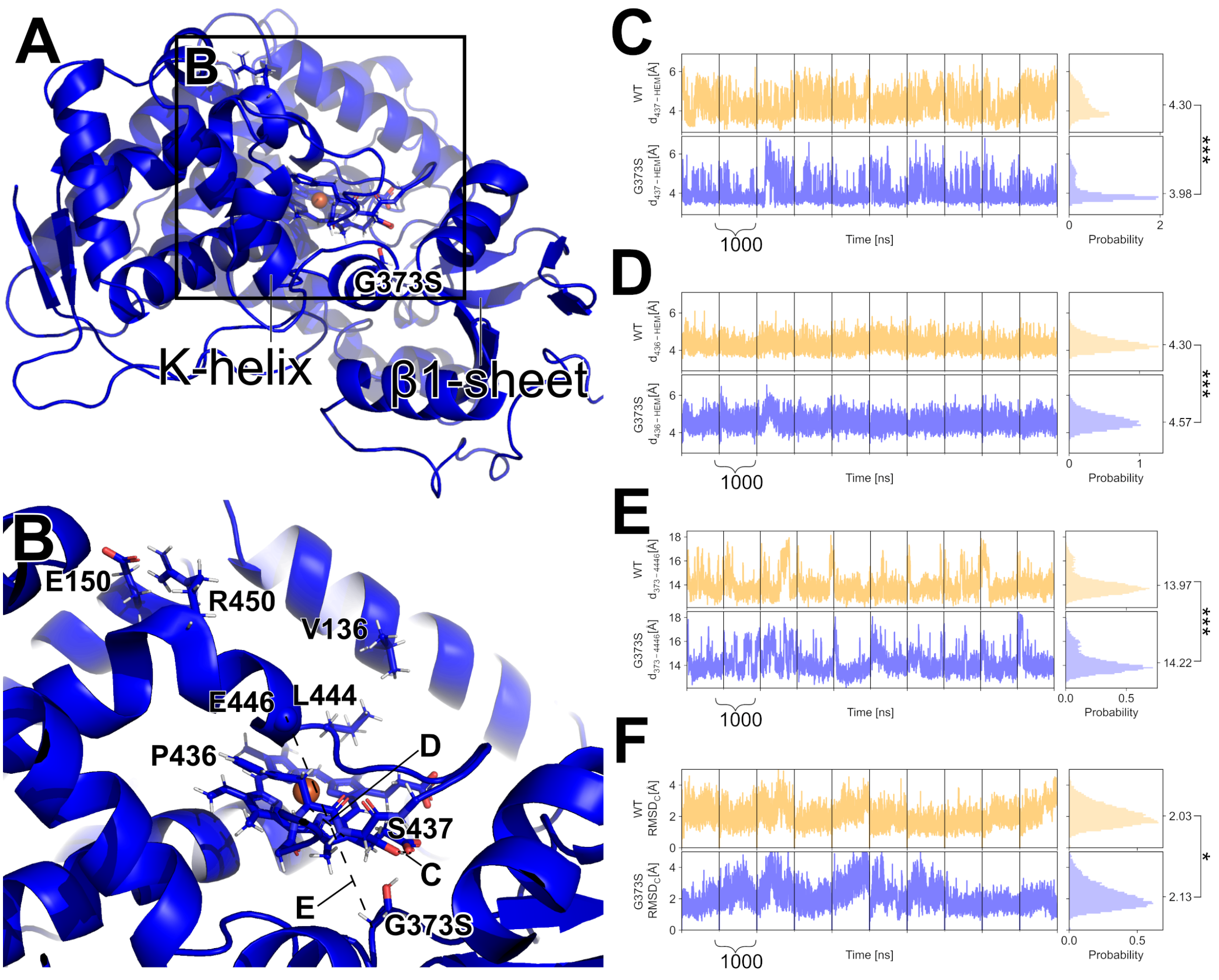


**Figure S11:** Effect of the G373S mutation on CYP2D6. **A:** The mutation in G373S is located in the K-K'-loop. The heme ring system is shown as sticks, while the heme iron and important $C_\alpha$ atoms are shown as spheres. **B:** The OH group in the variant stabilizes the position of the close-by carbonyl group in the heme moiety. This stabilizes the position of S437, which lifts the K-L-loop relative to the heme. This lifting leads to a higher distance between the $C_\alpha$ atom of residue G373/S373 (see panel E) and the $C_\alpha$ atom of residue E446 because of the interactions between P436 and the backbone of E446. This also increases the hydrophobic interaction between L444 and V136. Dashed lines indicate measured distances. **C:** The interaction between S437-OH and the heme carboxy group of ring D is significantly increased in G373S. **D:** The $C_\alpha$ atom of P436 shows an increased distance to the aromatic carbon 17 in ring D of heme. **E:** The distance between the $C_\alpha$ atom of residue G373/S373 and the $C_\alpha$ atom of residue E446 is significantly increased ($p < 0.01$, two-sided $t$-test). C-E: Left: time series of the distance, black vertical lines separate values per trajectory; right: aggregate probability distributions. **F:** The $C_\alpha$-RMSD of the C-helix is significantly increased in G373S. Left: time series of the $C_\alpha$-RMSD, values per trajectory are separated by black vertical lines; right: aggregate probability distributions. Statistical analysis was performed using the two-sided t-test (* $p < 0.01$;** $p < 0.001$; *** $p < 0.0001$; n.s.: $p > 0.01$).

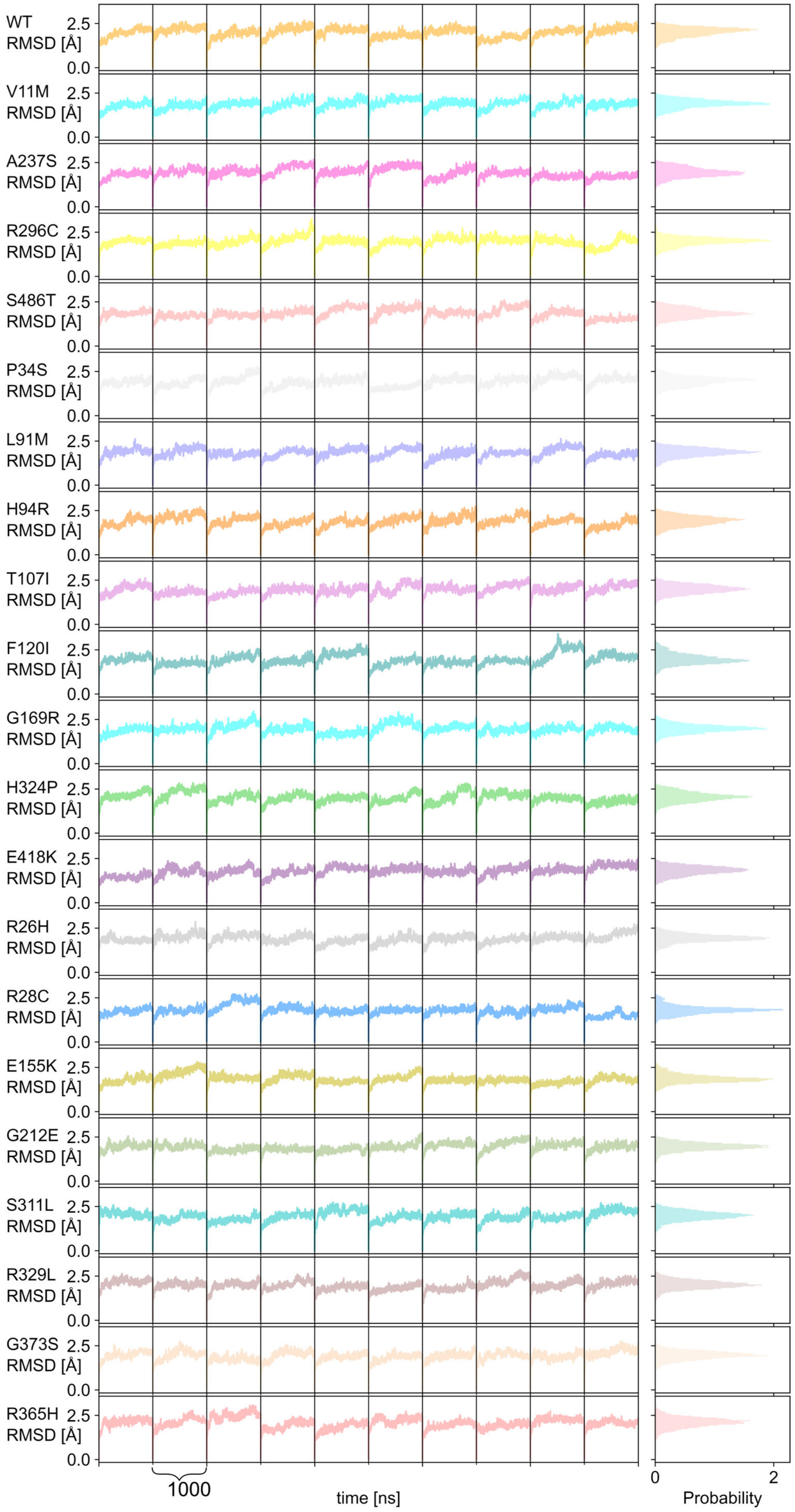
WT
V11M
A237S
R296C
S486T
P34S
L91M
H94R
T107I
F120I
G169R
H324P
E418K
R26H
R28C
E155K
G212E
S311L
R329L
G373S
R365H
RMSD [Å]
2.5
0.0
1000
time [ns]
0
2
Probability

**Figure S12: Root-mean-square deviation (RMSD) of the backbone of the CYP2D6 wild type and variants along ten MD trajectories each and aggregate probability distributions.** The RMSD is calculated with respect to the structure in the first frame after equilibration. The first 50 residues were not considered in the RMSD calculations because the TM-helix can move independently in the membrane.

### Supplemental References

1. Rathi, P. C.; Mulnaes, D.; Gohlke, H., VisualCNA: a GUI for interactive constraint network analysis and protein engineering for improving thermostability. *Bioinformatics* **2015**, 31, 2394-6.
2. Walt, S. v. d.; Colbert, S. C.; Varoquaux, G., The NumPy Array: A Structure for Efficient Numerical Computation. *Computing in Science & Engineering* **2011**, 13, 22-30.
3. Virtanen, P.; Gommers, R.; Oliphant, T. E.; Haberland, M.; Reddy, T.; Cournapeau, D.; Burovski, E.; Peterson, P.; Weckesser, W.; Bright, J.; van der Walt, S. J.; Brett, M.; Wilson, J.; Millman, K. J.; Mayorov, N.; Nelson, A. R. J.; Jones, E.; Kern, R.; Larson, E.; Carey, C. J.; Polat, İ.; Feng, Y.; Moore, E. W.; VanderPlas, J.; Laxalde, D.; Perktold, J.; Cimrman, R.; Henriksen, I.; Quintero, E. A.; Harris, C. R.; Archibald, A. M.; Ribeiro, A. H.; Pedregosa, F.; van Mulbregt, P.; Vijaykumar, A.; Bardelli, A. P.; Rothberg, A.; Hilboll, A.; Kloeckner, A.; Scopatz, A.; Lee, A.; Rokem, A.; Woods, C. N.; Fulton, C.; Masson, C.; Häggström, C.; Fitzgerald, C.; Nicholson, D. A.; Hagen, D. R.; Pasechnik, D. V.; Olivetti, E.; Martin, E.; Wieser, E.; Silva, F.; Lenders, F.; Wilhelm, F.; Young, G.; Price, G. A.; Ingold, G.-L.; Allen, G. E.; Lee, G. R.; Audren, H.; Probst, I.; Dietrich, J. P.; Silterra, J.; Webber, J. T.; Slavič, J.; Nothman, J.; Buchner, J.; Kulick, J.; Schönberger, J. L.; de Miranda Cardoso, J. V.; Reimer, J.; Harrington, J.; Rodríguez, J. L. C.; Nunez-Iglesias, J.; Kuczynski, J.; Tritz, K.; Thoma, M.; Newville, M.; Kümmerer, M.; Bolingbroke, M.; Tartre, M.; Pak, M.; Smith, N. J.; Nowaczyk, N.; Shebanov, N.; Pavlyk, O.; Brodtkorb, P. A.; Lee, P.; McGibbon, R. T.; Feldbauer, R.; Lewis, S.; Tygier, S.; Sievert, S.; Vigna, S.; Peterson, S.; More, S.; Pudlik, T.; Oshima, T.; Pingel, T. J.; Robitaille, T. P.; Spura, T.; Jones, T. R.; Cera, T.; Leslie, T.; Zito, T.; Krauss, T.; Upadhyay, U.; Halchenko, Y. O.; Vázquez-Baeza, Y.; SciPy, C., SciPy 1.0: fundamental algorithms for scientific computing in Python. *Nat. Methods* **2020**, 17, 261-272.
4. Roe, D. R.; Cheatham, T. E., PTRAJ and CPPTRAJ: Software for Processing and Analysis of Molecular Dynamics Trajectory Data. *J. Chem. Theory Comput.* **2013**, 9, 3084-3095.
5. Homeyer, N.; Gohlke, H., FEW: A workflow tool for free energy calculations of ligand binding. *J. Comput. Chem.* **2013**, 34, 965-973.
6. Åqvist, J.; Luzhkov, V. B.; Brandsdal, B. O., Ligand Binding Affinities from MD Simulations. *Acc. Chem. Res.* **2002**, 35, 358-365.
7. Brandsdal, B. O.; Österberg, F.; Almlöf, M.; Feierberg, I.; Luzhkov, V. B.; Åqvist, J. Free Energy Calculations and Ligand Binding. In *Adv. Protein Chem.*; Academic Press: 2003; Vol. 66, pp 123-158.
8. Biedermannova, L.; E. Riley, K.; Berka, K.; Hobza, P.; Vondrasek, J., Another role of proline: stabilization interactions in proteins and protein complexes concerning proline and tryptophane. *Phys. Chem. Chem. Phys.* **2008**, 10, 6350-6359.
